# Evolutionary diversification of invertase paralogs couples carbon metabolism and sexual reproduction in fission yeasts

**DOI:** 10.64898/2026.04.13.718246

**Authors:** Ambre Noly, Céline Faux, Maria Shaldaeva, Louison Bois-Naegelin, Philippe Fort, Dominique Helmlinger

## Abstract

Dynamic patterns of gene gain and loss play a major role in the diversification of eukaryotes, reflecting adaptation to a broad range of ecological contexts. Reconstructing the evolutionary history of genes provides a powerful framework for understanding how functional innovation shapes life-history traits. Here we report a comprehensive analysis of gene gain and loss across the fission yeast clade, whose evolutionary trajectory remains elusive. Reductive evolution of metabolic genes is a major contributor to species diversification, as observed in other fungal taxa. Notably, we uncovered an evolutionary scenario in which an ancestral gene duplication was followed by lineage-specific loss of one or the other paralog, except in *S. pombe*, which retained both. We demonstrate that these paralogs encode catalytically-active invertases, named Inv1 and Inv2, with distinct enzymatic properties, localization, regulation, and physiological roles. Inv1 is a secreted enzyme subject to glucose catabolite repression and is the sole invertase required for sucrose assimilation, resembling canonical yeast invertases. In contrast, Inv2 is intracellular, constitutively expressed, and required for inducing sexual differentiation in response to nutrient availability. Overall, these findings reveal an unexpected role for carbon metabolism in modulating the haploid-diploid cycle of fission yeasts, suggesting that diversification of core metabolic functions may contribute to adaptation to environments with distinct sugar compositions.

**Significance Statement:** Although *Schizosaccharomyces pombe* is a prime model organism, little is known about the evolutionary history and ecological adaptations of the broader fission yeast clade. By analyzing patterns of gene gain and loss across fission yeasts, this study identifies metabolic genes as major contributors to genomic diversification and uncovers an unusual case of ancestral gene duplication followed by reciprocal paralog loss. We show that *S. pombe* uniquely retained both paralogs, which encode distinct invertases that have diverged in regulation, localization, and physiological roles. Functional specialization of these enzymes links carbon metabolism to both nutrient assimilation and the control of sexual differentiation. Together, these findings connect genome evolution to metabolic specialization and life-cycle regulation, providing new insights into the ecological drivers of genome function and evolution in unicellular eukaryotes.

## Introduction

Yeasts occupy a wide range of ecological niches and exhibit highly divergent life history traits. For example, distinct strategies for alternating between asexual and sexual reproduction evolved across the fungal tree of life. Whereas *Saccharomyces* yeasts mate constitutively and require a non-fermentable carbon source to undergo meiosis, *Schizosaccharomyces* species are haplobiontic organisms in which mating is a specific response to nitrogen starvation. These evolutionary adaptations have a profound impact on genome organization (Leducq 2014). In this context, comparative genomics emerged as a powerful approach to identify the genetic determinants of adaptation to contrasting environments. Seminal work focusing on the Saccharomycotina subphylum revealed that ploidy changes, gene duplication, horizontal gene transfer (HGT), and introgression played important roles in shaping yeast genome evolution (Marsit et al. 2017). Notably, as in other eukaryotic lineages, gene gain and loss are major drivers of evolutionary and ecological transitions in yeasts, facilitating the emergence of novel traits and the colonization of diverse ecological niches (Wu et al. 2022; Domazet-Lošo et al. 2024).

Metabolic genes and pathways are frequently remodeled during genome evolution as they operate at the interface with the environment, thereby providing key insights into the diversity of trophic strategies and ecological interactions (Shen et al. 2018; Opulente et al. 2024). In yeasts, metabolic diversity ranges from generalist species, capable of utilizing a wide array of substrates, to specialists restricted to only one or a few. Carbon metabolism, in particular, is a highly variable trait across species and strains, and a large repertoire of enzymes, transporters, and regulatory factors evolved to support growth and survival on a range of different carbon sources.

The ability of yeasts to assimilate sucrose, the most abundant sugar produced by land plants, illustrates how a metabolic trait diversified across budding yeast species. Sucrose fermentation and utilization positively correlate with a generalist lifestyle and presence in flowers, fruits, fermented substrates, and floricolous insects (Opulente et al. 2018; Gonçalves et al. 2020; Opulente et al. 2024). For instance, most *Wickerhamiella* and *Starmerella* species thrive in floral niches and, yet, independently evolved distinct sucrose assimilation strategies (Gonçalves et al. 2022). Sucrose is consumed either extracellularly by a bacterial invertase acquired though HGT, or intracellularly using broad substrate-range α-glucoside/H^+^ symporters and α-glucosidases with complex evolutionary histories (Gonçalves et al. 2022). In this clade, alcoholic fermentation also shows dynamic evolutionary remodeling, including loss and subsequent reacquisition, reflecting divergent ecological adaptations (Gonçalves et al. 2020).

Sucrose is typically hydrolyzed by invertases, or β-fructofuranosidases, which belong to the glycoside hydrolase family GH32 (Drula et al. 2022). These enzymes specifically catalyze the hydrolysis of the α-1,2-glycosidic bond to release glucose and fructose. Invertase is among the first discovered enzymatic activities, identified in yeast during the mid-19th century, and its characterization established a paradigm for enzymology, protein synthesis, and glycoprotein secretion. In particular, Leonor Michaelis and Maud Menten used invertase to develop their foundational model of enzyme-catalyzed reactions (Johnson and Goody 2011). GH32 family invertases are found across the tree of life and widely distributed in plants, fungi, protists, and bacteria, but only present in a few invertebrate phyla (Parrent et al. 2009; Manoochehri et al. 2020; Cheng et al. 2024). Our knowledge of eukaryotic invertase function and regulation comes mostly from studies performed in plants, as sucrose plays a central role in their metabolism and physiology (Ruan 2014). Plants possess up to three distinct invertases with different biochemical properties, subcellular localization, and functions. Invertases with an acidic optimum pH are either extracellular or vacuolar and play key roles in plant growth, development, and stress responses. In contrast, cytosolic invertases are neutral or alkaline enzymes with only limited information available on their physiological roles.

Comparatively less is known about fungal invertases, except in the budding yeast *Saccharomyces cerevisiae*. In this species, six genes belonging to the *SUC* family encode an invertase enzyme. These genes are differentially present depending on the strain, with only *SUC2* being ubiquitous across strains that ferment sucrose (Mortimer and Hawthorne 1966). Suc2 enzymatic activity, oligomeric structure, glycosylation, secretion mechanisms, and expression regulation have been extensively characterized (Neumann and Lampen 1967; Bowski et al. 1971; Perlman and Halvorson 1981; Carlson and Botstein 1982; Carlson et al. 1983; Esmon et al. 1987; Reddy et al. 1988; Schülke and Schmid 1988; Ziegler et al. 1988; Reddy and Maley 1990; Sainz-Polo et al. 2013). Although *SUC2* is solely responsible for sucrose assimilation, early studies identified two invertase activities in *S. cerevisiae* (Gascón et al. 1968; Mwesigye and Barford 1996). Elegant genetic and molecular analyses demonstrated that *S. cerevisiae* produces two forms of invertase (Perlman and Halvorson 1981; Carlson and Botstein 1982; Carlson et al. 1983). One isoform is glycosylated, secreted and anchored to the cell wall, while the other is transcribed from an alternative promoter into a shorter isoform that lacks the N-terminal signal peptide. The latter isoform is thus non-glycosylated and retained in the cytoplasm. In addition, the expression of the extracellular enzyme is under glucose catabolite repression and is essential to use sucrose as a carbon source, while the intracellular isoform is constitutively expressed, at low levels, with no clear function described to date. Therefore, in *S. cerevisiae*, one gene encodes two invertase isoforms with distinct localization and regulation.

Beyond this case, only few additional fungal species possess distinct invertases (Goosen et al. 2007). The filamentous fungus *Aspergillus niger* has three invertase paralogs, including *sucB*, which encodes an intracellular enzyme that is not essential for sucrose catabolism (Goosen et al. 2007). Instead, *sucB* deletion accelerates sporulation in the presence of various carbon sources, suggesting that, similar to plants, certain fungi may metabolize sucrose within the cytosol to regulate physiological processes other than anabolic growth. In the fission yeast *Schizosaccharomyces pombe*, two distinct invertase activities were reported (Mitchison et al. 1969; Moreno et al. 1985). Although their catalytic properties differ between basal and derepressed conditions, both activities localize to the cell wall and are associated with a glycosylated polypeptide, unlike in *S. cerevisiae* (Mitchison et al. 1969; Moreno et al. 1985; Moreno et al. 1990a). However, only a single gene, *inv1*, was isolated (Tanaka et al. 1998). This gene is under glucose-catabolite repression and *inv1*Δ mutants lack detectable invertase activity, yet surprisingly, exhibit residual growth on sucrose when respiration is inhibited (Tanaka et al. 1998; Reinders and Ward 2001; Ahn et al. 2012). Altogether, these observations support the hypothesis that *S. pombe* possesses an additional invertase, whose activity depends on metabolic conditions, although whether this activity derives from a paralogous gene or an alternative isoform remains elusive.

Here we report a comprehensive analysis of gene gain and loss across the genomes of fission yeast species. We show that *S. pombe* possesses two distinct invertase genes, *inv1* and *inv2,* whereas other branches of the fission yeast clade lost either paralog. We demonstrate that both genes encode sucrose-hydrolyzing enzymes, although with distinct catalytic properties. Inv1 is glycosylated, secreted, under glucose-catabolite repression, and the only invertase essential for growth on sucrose, analogous to *S. cerevisiae* Suc2. In contrast, Inv2 is retained in the cytosol, expressed constitutively, and required to coordinate sexual reproduction with carbon availability, resembling *A. niger* SucB. Remarkably, these distinct functions are conserved across fission yeasts, spanning over 200 million years (Myr) of evolution. Together, our evolutionary and molecular analyses illustrate how gene duplication enables the evolutionary co-option of a metabolic enzyme for the regulation of an important life-history transition.

## Results

### Reconstruction of gene gain and loss history in the fission yeast clade

To date, the *Schizosaccharomyces* genus comprises seven species, including *S. japonicus* (Sj), *S. versatilis* (Sv), *S. pombe* (Sp), *S. cryophilus* (Sc), *S. octosporus* (So), *S. lindneri* (Sl), and *S. osmophilus* (Ss) (Figure 1A). Fission yeasts occur only sporadically within natural microbial communities, suggesting colonization of specific ecological niches and adaptation to highly specialized environments (Jeffares et al. 2015; Brysch-Herzberg et al. 2022). For example, bee food provisions, such as honey, appear as the primary habitats of *S. pombe*, *S. octosporus*, and *S. osmophilus*, while *S. japonicus* is mainly found in forest substrates and fruits (Brysch-Herzberg et al. 2022). We therefore investigated whether fission yeast genomes harbor distinctive genetic features, focusing on gene gains and losses, which are major drivers of adaptive evolution (Qian and Zhang 2014; Helsen et al. 2020).

**Figure 1:**
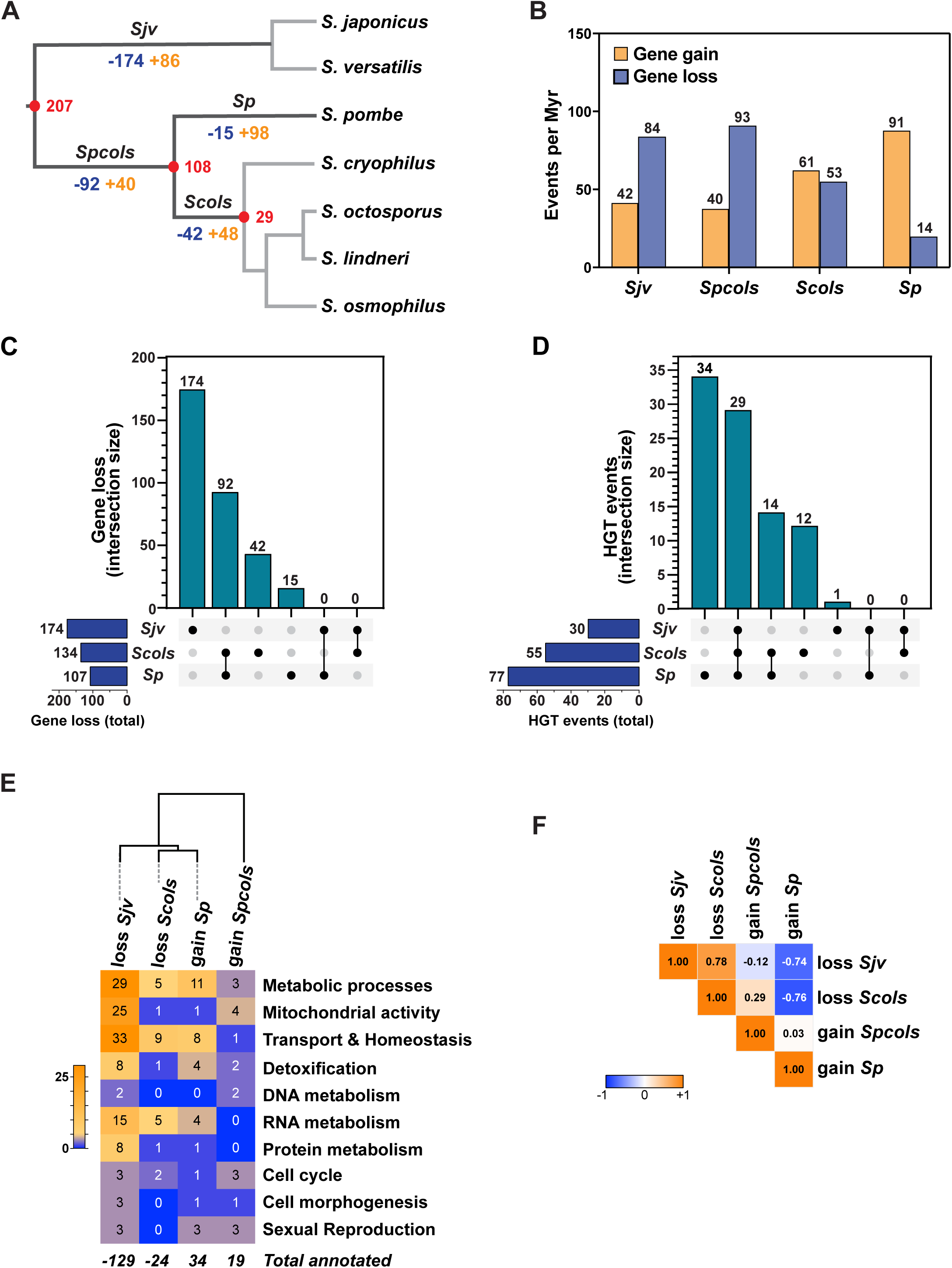
Gene and ontology evolutionary dynamics across the fission yeast clade. (A) Phylogenetic tree of fission yeast species based on (Brysch-Herzberg et al. 2024; Etherington et al. 2024). Shown are the number of genes gained (orange) or lost (blue) along each lineage. The *Sjv* branch, comprising *S. japonicus* and *S. versatilis* diverged from the *Spcols* lineage, which subsequently split into the *Sp* branch, with *S. pombe* as the sole representative, and the *Scols* branch, which includes *S. cryophilus, S. octosporus*, *S. lindneri* and *S. osmophilus*. Red nodes indicate estimated divergence times in Myr. (B) Bar graph representing the number of gene gain and loss events divided by the divergence time for each indicated lineage, and expressed per 100 Myr. (C) UpSet plot visualizing the total number of HGT events in each lineage (bottom left bar plot) and the intersection of events to reveal shared and specific acquisitions (top bar plot and matrix). (D) UpSet plot visualizing the total number of gene loss events in each lineage (bottom left bar plot) and the intersection of events to reveal shared and specific losses (top bar plot and matrix). (E) Hierarchical clustering of functionally annotated genes that were either lost in *Sjv*, lost in *Scols*, gained in *Sp*, or gained in *Spcols*. Gene Ontology Biological Process terms were grouped into ten categories (see Table S2) before clustering. Shown are the absolute number of gain and loss events for each category using a sequential colour scale. (F) Heat map representation of the correlation between ontology gains and losses across the lineages shown in (D). Shown are Pearson correlation coefficients using a sequential colour scale.

Approximately 207 million years ago (Mya), an initial speciation event gave rise to two lineages. One branch led to *S. japonicus* and *S. versatilis* (*Sjv*), which separated between 13 and 25 Mya (Brysch-Herzberg et al. 2024; Etherington et al. 2024). The availability of two genomes ensures a more accurate characterization of gain and loss events in this lineage. The other phylogenetic branch is ancestral to the five other species, *S. pombe*, *S. cryophilus*, *S. octosporus*, *S. lindneri*, and *S. osmophilus* (*Spcols*). Around 100 Myr later, *S. pombe* (*Sp*) diverged from the branch that contains the four remaining species, *S. cryophilus*, *S. octosporus*, *S. lindneri*, and *S. osmophilus* (*Scols*), from which *S. cryophilus* separated first, about 29 Mya (Figure 1A). To investigate gene gain and loss events associated with the diversification of Schizosaccharomycetes, we focused on the *Sjv*, *Spcols*, *Sp*, and *Scols* branches, which span 207, 99, 108, and 79 Myr of evolution, respectively (Figure 1A). For this, we leveraged our recent comprehensive orthology analysis of all fission yeast protein-coding genes ((Jia et al. 2026) and Table S1). This resource integrates high-quality genome assemblies and gene annotations to provide robust orthology relationships between the seven fission yeast species. We have now identified a total of 5,414 distinct orthology relationships that exhibit either universal or patchy distributions. The latter represents about 18% of all protein-coding genes across the seven genomes, and here we sought to identify in which case this pattern can be explained by either a gain or a loss in each species and lineage.

The apparent absence of a gene in a given species can reflect either lineage-specific loss or, alternatively, a gain in other lineages, depending on the presence of homologs in more distantly related taxa. To root the phylogenetic analysis of all fission yeast genes with a patchy distribution, we performed reciprocal BLASTP and TBLASTN analyses, first against Taphrinomycotina, then against other fungal and bacterial genome sequences (Table S1). This strategy allowed reconstructing the history of gene gain and loss events across major node transitions within the clade, including HGT events (Figure 1A-D, Tables S1 and S2).

The evolutionary dynamics of gene content was broadly proportional to divergence times, with the notable exception of *S. pombe* (Figure 1A,B). In the *Sjv* lineage, we detected 86 gains and 174 losses over 207 Myr, corresponding to rates of 42 gains and 84 losses per 100 Myr. Gene gains and losses occurred at similar rates in the ancestral *Spcols* lineage, with 40 gains and 93 losses over the same time frame. Following the speciation of *S. pombe*, the rate of gene gain increased in the *Scols* lineage and was even higher in *Sp*, with 61 and 91 gains per 100 Myr, respectively. Conversely, both lineages showed lower rates of gene losses, particularly *Sp*, with 53 and only 14 events per Myr inferred in *Scols* and *Sp*, respectively (Figure 1B). Finally, we found no evidence of convergent gene loss between *Sjv* and either the *Sp* or *Scols* lineages (Figure 1C).

These observations indicate that *S. pombe* lost relatively fewer genes than the other species, consistent with its isolation from a broader range of substrates (Brysch-Herzberg et al. 2022), while acquiring a relatively high number of genes compared with the other lineages (Figure 1C,D). The number of inferred gene gains in *S. pombe* may be inflated by the fact that it is the sole representative of its subclade. However, to mitigate this potential bias, we retained only genes shared with the most divergent *S. pombe* isolate identified to date (Jia et al. 2026) and excluded 93 genes annotated as dubious in PomBase (Carme et al. 2026). Notably, we found that horizontal transfer of prokaryotic genes contributes substantially to these gains (Figure 1D), consistent with a previous study (Rhind et al. 2011). By comparing HGT events across the three main lineages, we identified a total of 90 events, of which 29 occurred prior to the radiation of the fission yeast clade. Subsequently, only one HGT was detected specifically in the *Sjv* lineage, whereas 14 prokaryotic genes were acquired along the *Spcols* branch, followed by 34 events specific to *Sp* and 12 specific to *Scols*. Thus, over a comparable evolutionary timescale, the five *Spcols* species acquired substantially more genes by HGT than the *Sjv* species, which might reflect differences in their environmental distribution and ecological niches.

### Distribution of gene gains and losses within gene ontology categories

In contrast to the other fission yeasts, *S. pombe* is a prime model organism, with a considerable wealth of information on its genetics, cell biology, and physiology. Consequently, gene gains and losses directly involving *S. pombe* orthologs can provide valuable biological information, in particular because of their comprehensive curation in the PomBase database and reliable Gene Ontology (GO) annotations (The Gene Ontology Consortium et al. 2023; Carme et al. 2026). We thus extracted from PomBase GO Biological Process (BP) terms associated with each *S. pombe* accession of gained and lost genes, annotating a total of 206 genes (Table S2). These include 129 genes lost specifically in *Sjv*, 24 in *Scols*, while 19 gene gains were annotated for *Spcols* and 34 for *Sp*. These associations distributed into 40 distinct BP terms, which we grouped into 10 main categories of related terms before hierarchical clustering analysis (Figure 1E and Table S2). Within each lineage, the distributions of gains and losses differ from the null hypothesis (*P*-values < 0.0001), except for the losses in the *Scols* branch (*P*-value = 0.19) (Table S2), suggesting that some functions appear specifically affected in *Sjv*, *Spcols*, and *Sp*. Notably, about 20% of annotated gene losses in *Sjv* show mitochondria related functions (Figure 1D), in agreement with the observed loss of respiration in *S. japonicus* (Kaino et al. 2018; Alam et al. 2023), validating the physiological relevance of our comparative ontology analysis.

We found that more than half of annotated *Sjv* and *Scols* losses involve factors contributing to metabolic processes or membrane and organelle transport (Figure 1E and Table S2). Interestingly, *S. pombe* speciation was accompanied by gains of genes involved in the same ontology categories. We also noticed the gain of three genes involved in meiotic progression and spore morphogenesis, including the spore wall structural component *isp3* (Fukunishi et al. 2014; Tahara et al. 2020). Given the importance of Isp3 in *S. pombe* spore resistance, it is likely that fission yeast species differ in spore morphology and stress tolerance, possibly related to adaption to different habitats. In contrast, the *Spcols* lineage gained genes distributed more homogenously across our ontology categories (Figure 1E and Table S2). These include genes with well-characterized roles in promoting meiosis, such as *mei3* (Vještica et al. 2021), and chromosome segregation, such as *nks1* (Buttrick et al. 2011; Chen et al. 2011), suggesting that these species evolved distinct regulatory mechanisms for the mitotic and meiotic cell cycles.

Finally, direct comparison of ontology gains and losses between lineages confirmed that *Sjv* and *Scols* appear to have lost overlapping functions, whereas *Sp* gained factors involved in the same processes, as indicated by the strong correlation between the distributions of their categories (Figure 1F and Table S2). These correlation coefficients are even higher when the ‘Mitochondrial activity’ category, specifically enriched in the *Sjv* branch, was omitted (Table S2). Indeed, although distinct genes are affected, nutrient transporters and metabolic factors were independently lost along the *Sjv* and *Scols* lineages but gained during the evolution of *Sp* (Figure 1E and Table S2). For example, the *Scols* lineage lost *pdc201*, a pyruvate decarboxylase contributing to long-term survival following exponential growth (Kim et al. 2014) and the sucrose-assimilating *inv1* invertase (Tanaka et al. 1998). In parallel, genes predicted to contribute to disaccharide assimilation, including the putative invertase *inv2* and the putative maltase *mal1* were lost from the *Sjv* branch.

In conclusion, the *Sjv* branch shows the strongest functional divergence, having lost many genes encoding factors involved in mitochondrial activity, as well as carbon, nitrogen, amino-acid, and nucleobase transport and metabolism. In contrast, *S. pombe* lost fewer genes while acquiring new functions, including through HGT.

### The fission yeast clade evolved two invertases

The presence of two paralogous genes in *S. pombe*, *inv1* and *inv2*, together with their reciprocal loss in the other lineages, caught our attention because it represents a unique evolutionary pattern in this clade (Table S2). Both genes encode proteins annotated as enzymes from the glycoside hydrolase family 32 with β-fructofuranosidase activity (GH32, EC #3.2.1.26), better known as invertases, which hydrolyze sucrose into glucose and fructose (www.cazy.org) (Henrissat 1991). To examine the divergence of fission yeast invertases, we performed pairwise alignments of their amino-acid sequences from Schizosaccharomycetes and *S. cerevisiae*. Surprisingly, Inv1 and Inv2 share only marginally higher sequence identity with each other than either does with *S. cerevisiae* Suc2 (Figure S1A,B). This pattern suggests that Inv1 and Inv2 either diverged extensively following an early duplication in the Taphrinomycotina lineage or have distinct evolutionary origins.

We therefore sought to distinguish between these two possibilities. The evolutionary history of fungal GH32 genes exhibits a complex pattern, comprising lineages with putative invertases, inulinases, and fructosyltransferases that do not follow species phylogeny (Parrent et al. 2009). In addition, previous work reported horizontal transfer of GH32 genes from bacterial to eukaryotic species, including fungi (Danchin et al. 2016; Noon and Baum 2016; Dai et al. 2021; Kfoury et al. 2024). To account for this complexity, we identified Inv1 and Inv2 homologs across nearly one hundred representative fungal, plant, and bacterial species, and reconstructed their phylogenetic relationships using maximum-likelihood analyses. As observed for GH32 enzymes, fungal invertases are dispersed across the tree with a topology that differs markedly from the species phylogeny (Figure S1C). Instead, they form two major clusters separated by groups of bacterial and plant sequences. The largest group, Cluster I, contains genes encoding invertases from most Dikarya species, including fission yeast Inv1 and Inv2, and corresponds to a sublineage of extracellular invertases identified in a previous study of GH32 phylogeny (Group 1 in (Parrent et al. 2009)). This cluster also includes prokaryotic invertases from α-, β-, and γ-proteobacteria of the Verrumicrobiota and Pseudomonadota clades, which branch at the root of this cluster with strong bootstrap support. The second group, Cluster II, consists primarily of Basidiomycota and Pezizomycotina sequences, which form two distinct sublineages that again do not follow the species phylogeny. This cluster also contains the three invertase paralogs from *A. niger*. These two groups correspond to those identified in earlier analyses of GH32 phylogeny and comprise both extracellular and intracellular invertases, including enzymes with experimentally demonstrated high fructosyltransferase activity (Group 7-8 in (Parrent et al. 2009) and VIb in (Alméciga-Díaz et al. 2011)).

To position Inv1 and Inv2 more precisely, we re-examined the phylogeny of invertases from Cluster I using Verrucomicrobiota sequences as an outgroup (Figure 2A). Both maximum-likelihood and Bayesian analyses confirmed the discrepancy between invertase and species phylogenies. Most internal nodes were weakly supported, possibly reflecting multiple and ancient acquisition of prokaryotic invertases through HGT by fungi. However, all fission yeast invertase sequences are connected with a statistically robust node, supporting the hypothesis that Inv1 and Inv2 originate from an ancestral gene duplication. We note that the branch leading to the three Inv1 sequences is much shorter than that leading to the five Inv2 sequences, suggesting that Inv2 may have evolved faster than Inv1 (Figure 2A). This difference could either reflect the ∼100 Myr difference in divergence times between the *Sjv* and *Scols* ancestors (Figure 1A) or, alternatively, result from asymmetric rates of paralog evolution following duplication, as previously described (Scannell and Wolfe 2008). However, estimating the nonsynonymous-to-synonymous substitution ratio (ω = *dN*/*dS*) for Inv1 and Inv2 is not feasible because the large evolutionary distance between *S. pombe* and the other fission yeast species results in saturation of synonymous substitutions (Fawcett et al. 2014; Harnqvist et al. 2021). We therefore could not test whether Inv1 and Inv2 have been subject to differential selective pressures.

**Figure 2:**
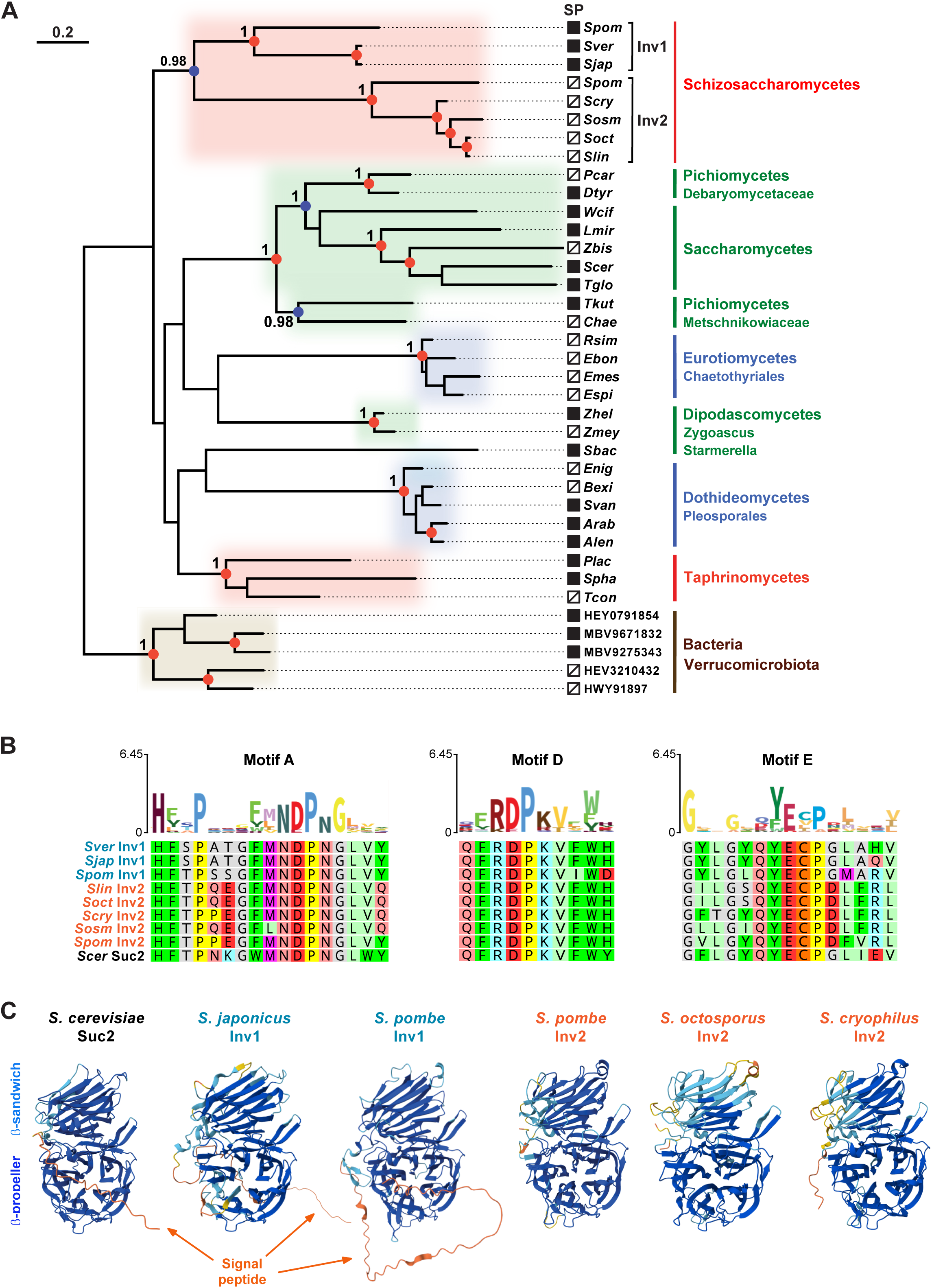
Inv2 is an ancient paralog of fungal extracellular invertases. (A) Phylogenetic relationship among the 32 Ascomycetes invertases belonging to Cluster I, with five Verrucomicrobiota sequences used as an outgroup (Figure S1C and Table S3). Sequences were aligned using MAFFT and filtered with BMGE (entropy cutoff = 0.7). The phylogenetic tree was inferred using bootstrapped maximum-likelihood analyses (n=100) and Bayesian inference (MrBayes). Red and blue circles indicate nodes supported by bootstrap values ≥90 and ≥70, respectively. Numbers above nodes indicate posterior probabilities ≥0.95. The scale bar refers to a phylogenetic distance of 0.2 amino-acid substitutions per site normalized to protein length. Filled black squares indicate proteins for which a signal peptide (SP) is predicted using SignalP 6.0 (Nielsen et al. 2024). Taxonomic classes and families are indicated on the right in large and small font sizes, respectively, and are color-coded according to the Ascomycota phylogeny with Taphrinomycotina in red, Saccharomycotina in green, and Pezizomycotina in blue. (B) Sequence logo representation of the A, D, and E catalytic motifs of Cluster I invertases and comparison with Schizosaccharomycetes invertases. Logos were generated from amino-acid sequence alignments of 59 proteins using the Skylign web tool. Shown are the regions corresponding to motifs A, D, and E, aligned with the corresponding motifs from all Schizosaccharomycetes invertase sequences. Asterisks indicate catalytic residues; underlined residues define sequence motifs specific to hydrolase *versus* fructosyltransferase activity, as described in (Trollope et al. 2015). (C) Predicted monomeric structures of invertases from, left to right, *S. cerevisiae* Suc2, *S. japonicus* Inv1, *S. pombe* Inv1, *S. pombe* Inv2, *S. octosporus* Inv2, and *S. cryophilus* Inv2 using AlphaFold via the EMBL-EBI server (Jumper et al. 2021). Structures are color-coded according to prediction confidence (pLDDT values). The N-terminal signal peptide, β-propeller and β-sandwich domains are annotated based on sequence alignments (Figure S2) and *S. cerevisiae* Suc2 crystal structure (Sainz-Polo et al. 2013).

We next investigated the conservation of known GH32 catalytic motifs in fission yeast invertases by generating sequence logos from multiple alignments of Cluster I invertases (Figure 2B). Comparison with the corresponding sequence motifs in Inv1 and Inv2 revealed conservation of all residues required for GH32 catalytic activity. These include the aspartate residues acting as catalytic nucleophiles in motifs A and D, and the glutamate residue functioning as a catalytic proton donor in motif E (Parrent et al. 2009). Closer examination of the sequence context surrounding these residues identified motifs specific to the hydrolase subgroup of GH32 enzymes, as opposed to those capable of fructo-oligosaccharide synthesis (Trollope et al. 2015). Likewise, AlphaFold-based prediction of Inv1 and Inv2 structures revealed the characteristic GH32 architecture, with a five-bladed β-propeller catalytic domain surrounding a negatively charged active-site cavity, appended to a β-sandwich domain (Figure 2C and S2). The predicted structure of Inv2 is nearly identical to that of Inv1 and to the crystal structure of the well-characterized *S. cerevisiae* Suc2 invertase (Sainz-Polo et al. 2013). However, in marked contrast to Inv1 and Suc2, all Inv2 orthologs lack a long, unstructured N-terminal region (Figure 2C and S2). Domain annotation and signal peptide predictions indicate that this region corresponds to a secretory signal, which is characteristic of extracellular invertases (Figure 2A).

In conclusion, we identified a second putative invertase of unknown function in fission yeast, Inv2, in addition to the previously described Inv1 enzyme (Tanaka et al. 1998). Phylogenetic analyses and sequence comparisons demonstrate that fission yeast Inv1 and Inv2 are ancient paralogs that retained key features of sucrose-hydrolyzing enzymes over at least 200 Myr of evolution, with the notable exception of secretion signal loss in Inv2.

### *S. pombe* Inv2 is a *bona fide* invertase

We next performed *in vitro* enzymatic assays using the endogenous protein affinity purified from *S. pombe* liquid cultures. For comparison, we purified Inv1 using the same C-terminal GFP epitope tag and procedure (Figure 3A). Total protein staining of Inv1 and Inv2 eluates showed that each enzyme is recovered with high purity and indicated that neither invertase interacts with other proteins. Sucrose hydrolysis was detected specifically in the presence of either Inv1 or Inv2, demonstrating that Inv2 can function as an invertase under these experimental conditions (Figure S3A). We then verified that Inv2 activity was measured within the linear range of the reaction by monitoring its activity over time (Figure S3B).

**Figure 3:**
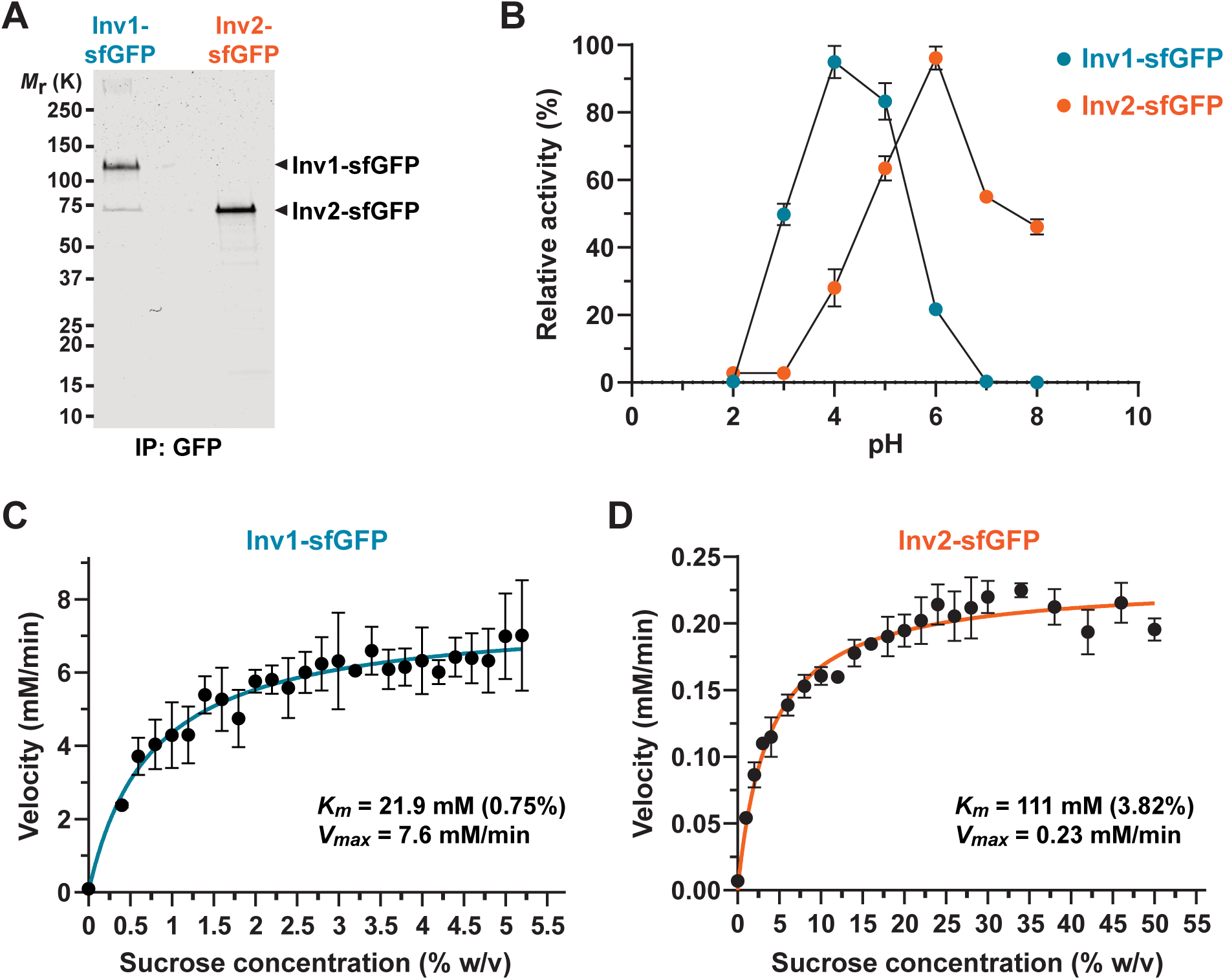
Sucrose-hydrolysing activity of *S. pombe* Inv2. (A) Coomassie blue staining of affinity purified Inv1-sfGFP and Inv2-sfGFP eluates. (B) Effect of pH on Inv1-sfGFP and Inv2-sfGFP invertase activities. Shown are enzymatic velocity values (mM/min) expressed as a percentage of the maximal measurement. Dots represent the mean of three independent experiments overlaid with the SD. (C,D) Enzyme kinetics of Inv1-sfGFP (C) and Inv2-sfGFP (D). Affinity purified Inv1-sfGFP and Inv2-sfGFP eluates were incubated with increasing sucrose concentrations, as indicated. Dots represent the mean of three independent experiments overlaid with the SD. Coloured lines represent the fit of a Michaelis-Menten nonlinear regression, from which the maximal velocity (*V_max_*) and the concentration of substrate required for the enzyme to achieve half *V_max_* (*K_m_*) were calculated.

We further characterized Inv1 and Inv2 catalytic properties by determining their optimal pH and found that Inv2 activity peaked at pH 6, while Inv1 reached maximal activity at a more acidic pH (Figure 3B), indicating different enzymatic behaviors. Using these parameters, we performed enzyme kinetics analyses by measuring glucose accumulation upon increasing concentrations of sucrose (Figure 3C,D). Non-linear regression fitting of each curve demonstrated that both Inv1 and Inv2 follow a Michaelis-Menten kinetics, but with distinct parameters. First, the sucrose concentration at which the reaction rate is half of the maximal velocity (*K_m_*) is about 3-fold higher for Inv2 (111 mM) compared to Inv1 (21.9 mM). Second, using comparable amounts of enzymes (Figure 3A), we found that Inv2 maximal velocity (*V_max_*) is about 30-fold less than that of Inv1, with 0.23 mM/min for Inv2 and 7.6 mM/min for Inv1. In conclusion, these results indicate that *S. pombe* Inv2 is a *bona fide* invertase, but has lower affinity and catalytic activity than Inv1. In contrast, *S. pombe* Inv1 exhibits enzymatic properties characteristic of canonical invertases, such as *S. cerevisiae* Suc2 (Nadeem et al. 2015).

### *S. pombe* invertase subcellular localization and expression regulation

Following up on our observation that Inv2 orthologs lack a secretion signal (Figure 2A,C and S2), we next examined the subcellular localization of *S. pombe* Inv2, as compared to Inv1, using the endogenous Inv1 and Inv2 GFP-tagged strains. Live confocal microscopy of *S. pombe* cultures revealed distinct localization of each enzyme as compared to background fluorescence levels (Figure 4A). Inv1 showed a strong signal at the periphery of cells, most likely external, as expected from the presence of a predicted signal peptide at its N-terminus (Figure 2A,C and S2). In marked contrast, Inv2 shows a diffuse intracellular localization throughout the cytoplasm and possibly the nucleus, consistent with the absence of a predicted secretion signal. Supporting these observations, we found that Inv1 and Inv2 display distinct electrophoretic migration pattern due to protein glycosylation (Figure 4B). Indeed, signal peptides target nascent polypeptides to the endoplasmic reticulum before glycosylation and secretion (Corsi and Schekman 1996). We observed that affinity purified Inv2 migrates at the expected size regardless of endoglycosidase treatment, while Inv1 apparent molecular weight shifts to the predicted size upon deglycosylation (Figure 4B).

**Figure 4:**
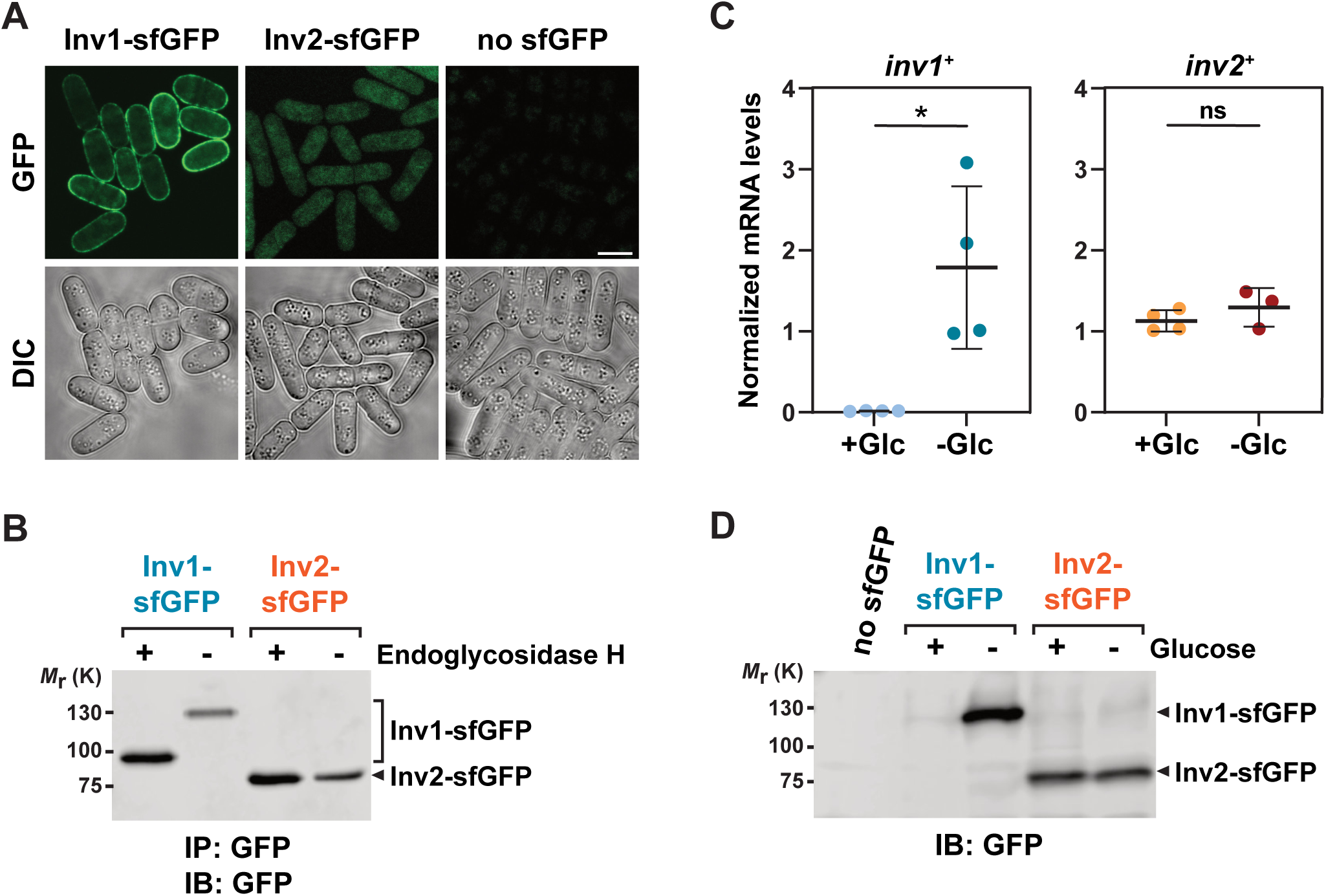
Unlike Inv1, Inv2 is intracellular, not under glucose-catabolite repression, and not glycosylated. (A) Cellular localization of *S. pombe* invertases. Live fluorescence microscopy (upper panels) of Inv1-sfGFP in low-glucose media (left) and Inv2-sfGFP in high-glucose media (middle). A ‘no-sfGFP’ control strain grown is included to show background fluorescence (right). Bottom panels show differential interference light microscopy images of the corresponding fields. Scale bar: 6 μm. (B) Anti-GFP Western-blot analysis of affinity purified Inv1-sfGFP and Inv2-sfGFP. Beads were treated (+) or not (-) with Endoglycosidase H to reveal Inv1 glycosylation. (C) Quantitative RT-PCR analysis of *inv1* and *inv2* mRNA levels in total RNA extracts from *S. pombe* grown to exponential phase in high- or low-glucose conditions (+Glc and -Glc, respectively). *snR92* served as a control for normalization across samples. Shown are the mean value of 3-4 independent biological replicates, overlaid with individual data points and the SD. Statistical significance was determined by unpaired, two-tailed t-tests with Welch’s correction. \**p* ≤ 0.05. (D) Anti-GFP Western-blot analysis of Inv1-sfGFP and Inv2-sfGFP levels in total protein extracts from *S. pombe* grown to exponential phase in high- (+) or low- (-) glucose conditions. A ‘no-sfGFP’ control strain grown in high-glucose medium is included to show antibody specificity.

We next tested whether Inv2 expression is under glucose-catabolite repression, as shown of *S. pombe* Inv1 (Tanaka et al. 1998; Ahn et al. 2012). Quantitative RT-PCR analyses demonstrated that glucose starvation induces *inv1* expression, whereas *inv2* mRNA levels remain constant (Figure 4C), in agreement with published transcriptome analyses (Rhind et al. 2011; Malecki et al. 2016; Vassiliadis et al. 2019). In agreement, Western blot analyses performed in the same conditions showed that GFP-tagged Inv1 is only detectable in *S. pombe* cells grown in low glucose conditions, unlike GFP-tagged Inv2, which is stably expressed (Figure 4D). We conclude that *S. pombe* Inv1 is secreted and under glucose-catabolite repression, whereas Inv2 is an internal invertase whose expression is not regulated by glucose availability.

### Roles of fission yeast invertases in sucrose assimilation

Our results so far demonstrate that *S. pombe* possesses two catalytically active invertases. However, Inv1 and Inv2 differ in their enzymatic properties, localization, and expression regulation, which may reflect adaptation to distinct metabolic cues and physiological contexts. To gain insights into their functions, we first tested sucrose assimilation using *S. pombe* deletion mutants. We found that *inv1*Δ strains exhibit a pronounced growth defect on sucrose media, whereas *inv2*Δ mutants have no observable phenotype relative to an isogenic wild-type (WT) strain (Figure 5A). Interestingly, *inv1*Δ mutants retain the ability to grow on media with sucrose as the sole carbon source, consistent with earlier reports (Tanaka et al. 1998; Reinders and Ward 2001). We thus hypothesized that Inv2 may partially compensate for the loss of Inv1. However, double *inv1*Δ *inv2*Δ deletion mutants show the same phenotype as single *inv1*Δ mutants, demonstrating that Inv2 is not required for sucrose assimilation in these conditions. Surprisingly, this observation suggests that *S. pombe* metabolizes sucrose even in absence of any invertases. We reasoned that sucrose hydrolysis by an external enzyme, such as Inv1, might enable glucose and fructose to diffuse and support growth of nearby colonies. Indeed, isolating strains by removing the agar abolished the growth of *inv1*Δ mutants on solid sucrose media (Figure 5B). Likewise, in liquid cultures, neither single *inv1*Δ nor double *inv1*Δ *inv2*Δ mutants proliferate in minimal media with sucrose. In contrast, *inv2*Δ deletion mutants grow exponentially and are indistinguishable from a WT control strain (Figure 5C,D). To test whether Inv2 is able to fulfil Inv1 function *in vivo*, we fused the Inv1 signal peptide sequence to the N-terminal end of Inv2 and placed this construct under a strong, inducible promoter (Lyu et al. 2024), to mimic *inv1* induction upon glucose starvation (Figure S4A,B). However, we observed that Inv2 remains cytoplasmic, possibly because it lacks glycosylation sites, and is unable to rescue the sucrose assimilation defect of *inv1*Δ mutants. Finally, we verified that the strain expressing endogenously sfGFP-tagged Inv1 used for enzymatic assays does not show any sucrose-assimilation defect, indicating that this version of Inv1 is fully functional (Figure S4C). This result is consistent with previous complementation experiments showing that a wild-type *inv1* copy restores sucrose assimilation in *inv1*Δ mutants (Tanaka et al. 1998). We conclude that Inv1 is the only invertase essential for sucrose assimilation in *S. pombe* grown in standard conditions.

**Figure 5:**
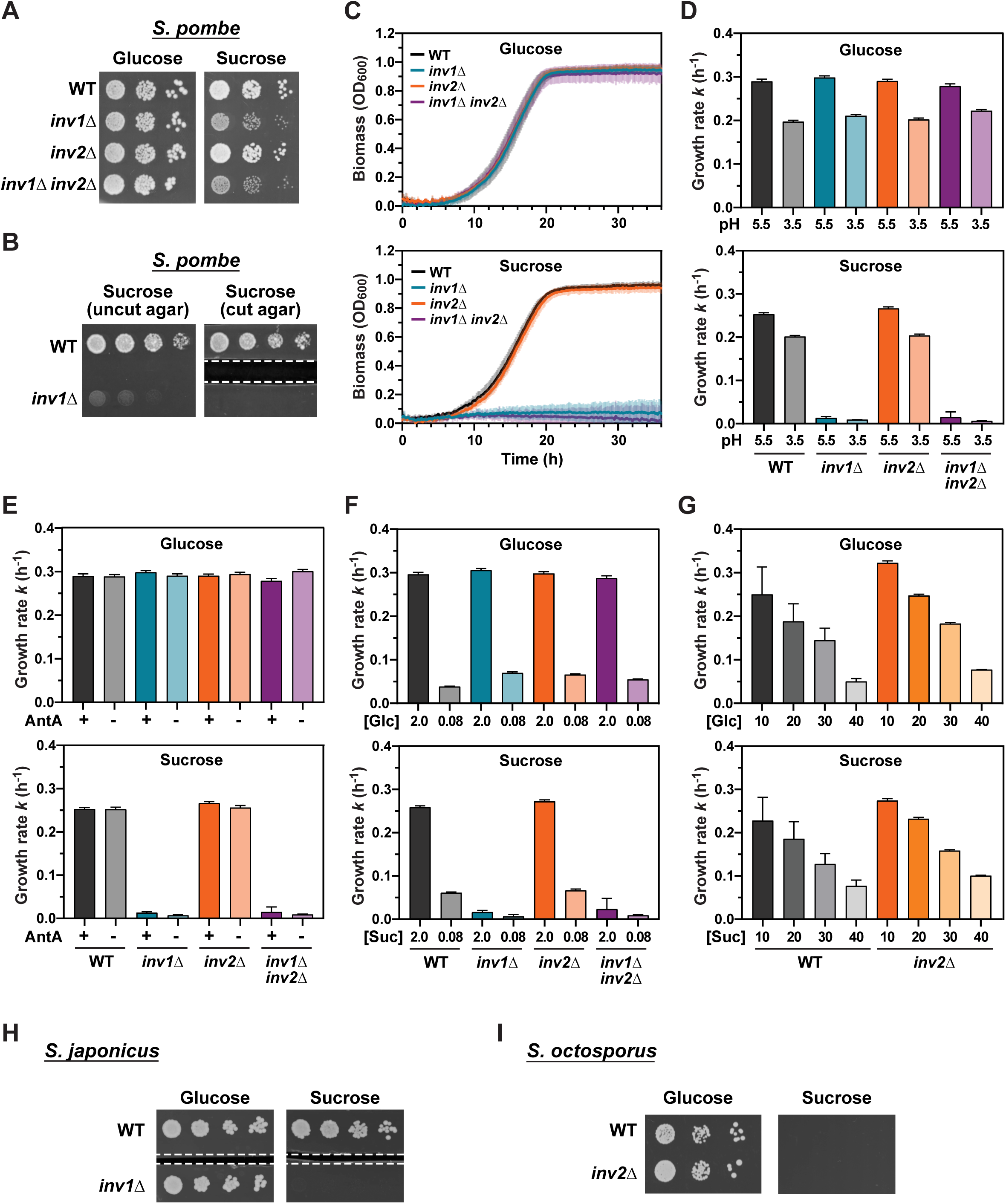
Unlike Inv1, Inv2 is not required for sucrose assimilation in fission yeasts. (A-G) Sucrose assimilation of *inv1*Δ, *inv2*Δ, and *inv1*Δ *inv2*Δ deletion mutants compared to isogenic WT controls in *S. pombe*. (A,B) Ten-fold serial dilutions of exponentially growing cells of the indicated genotypes were spotted in rich medium containing 3% glucose or sucrose and incubated at 32°C for 2 days. Shown are images that are representative of three independent experiments. White dashed lines in (B) represent agar cut out from the plates to prevent diffusion of nutrients. (C-G) Saturated cultures were diluted into liquid minimal media supplemented with 2% glucose or sucrose, unless indicated otherwise (F,G), and incubated at 32°C for 36 hours. Biomass was measured as a proxy of growth and monitored continuously by measuring absorbance (OD600) at 10 mins intervals. For quantitative comparisons between genotypes and conditions, growth rate values (*k*) were computed by fitting a modified logistic growth equation to proliferation curves and shown in (D-G). Shown are the mean value of 3 (F) or 4 (C-E,G) independent biological replicates overlaid with the SD. (H,I) Sucrose assimilation of *S. japonicus inv1*Δ (H) and *S. octosporus inv2*Δ (I) deletion mutants compared to isogenic WT controls. Ten-fold serial dilutions of exponentially growing cells of the indicated genotypes were spotted in rich medium containing 3% (H) or 30% (I) glucose and sucrose, and incubated at 30°C for 4 days. Shown are images that are representative of three independent experiments.

We then asked whether Inv2 contributes to sucrose catabolism in other metabolic contexts. For this, we repeated the liquid proliferation assays in different sucrose media, searching for a condition affecting the growth of *inv2*Δ single or *inv1*Δ *inv2*Δ double mutants. The intracellular localization of Inv2 (Figure 4A) prompted us to explore whether conditions favoring sucrose import might uncover the physiological role of Inv2. In *S. pombe*, Sut1 is the only characterized disaccharide transporter and mediates maltose and sucrose uptake via a proton-coupled symport mechanism (Reinders and Ward 2001). Since Sut1 transport activity increases at lower pH, we measured the growth of all deletion mutants in minimal media adjusted to pH 3.5, compared to normal pH, at 5.5 (Figure 5D). We observed that media acidification decreases the growth rate of all strains, in both glucose- and sucrose-containing media. However, at both pH, *inv2*Δ deletion mutants show no growth defects, as compared to a WT control, whereas both single *inv1*Δ and double *inv1*Δ *inv2*Δ mutants do not assimilate sucrose. Likewise, a *sut1*Δ deletion mutant did not show any growth defect in the presence of sucrose, regardless of pH (Figure S5A,B).

The products of sucrose hydrolysis, glucose and fructose, are catabolized to produce energy via either fermentation or respiration, prompting us to test whether Inv2 might be specifically required for either metabolic pathway. Supporting this hypothesis, Inv2 was specifically lost from *S. japonicus* (Figure 2A), which produces energy exclusively via fermentation (Kaino et al. 2018). Since *S. pombe* minimal media enables both fermentative and respiratory growth (Malecki et al. 2016), we first tested the role of Inv2 during fermentation using Antimycin A, which inhibits respiration by blocking mitochondrial electron transport at complex III (Figure 5E). We observed that Antimycin A has no effect on growth rates, regardless of strain genotype and carbon source. Conversely, as high glucose concentrations repress respiration in *S. pombe* (Takeda et al. 2015), we promoted respiratory growth by lowering the concentration of glucose and sucrose in the media from 2 to 0.08% (Figure 5F). Although lower carbon levels strongly reduce the growth rate of all strains in glucose and sucrose media, *inv2*Δ mutants remain able to use sucrose and to grow as WT controls, whereas both *inv1*Δ and *inv1*Δ *inv2*Δ mutants are unable to assimilate sucrose. We conclude that Inv1 is the only essential invertase whether *S. pombe* relies on fermentation or respiration for proliferation.

Sucrose breakdown by intracellular invertases is linked to osmotic regulation in plants (Liu, Pan, et al. 2025; Liu, Cheng, et al. 2025). In addition, the main habitats of fission yeast species that carry Inv2 are characterized by a high osmotic pressure, such as honey from honeybees and beebread from solitary bees, consistent with their ability to tolerate high glucose levels (Brysch-Herzberg et al. 2019; Brysch-Herzberg et al. 2022). These observations prompted us to test if Inv2 is required for growth in hyperosmotic conditions, by increasing the concentrations of glucose or sucrose in the media (Figure 5G). We observed that the growth rates of both *inv2*Δ mutant and control strains decrease as the levels of glucose and sucrose increase, but without any noticeable differences between genotypes, indicating that Inv2 is dispensable for osmotolerance in *S. pombe*.

Altogether, our results indicate that Inv1 is the only invertase essential for sucrose utilization in *S. pombe*, regardless of pH, metabolic conditions, or osmotic pressure and is thus the functional analogue of other yeast invertases, such as *S. cerevisiae* Suc2. Supporting this conclusion, we found that an *S. japonicus inv1*Δ deletion mutant is unable to grow on sucrose-containing media (Figure 5H), while wild-type *S. octosporus*, which naturally lacks Inv1, does not metabolize sucrose (Figure 5I). Finally, as observed in *S. pombe*, an *S. octosporus inv2*Δ deletion mutant shows no obvious fitness defect under standard growth conditions (Figure 5I).

### Roles of fission yeast Inv2 in sexual differentiation

Since Inv2 is not required for vegetative growth in the presence of sucrose or glucose, we next tested other carbon sources. Similarly, we found that Inv2 is dispensable for assimilation of maltose, trehalose, raffinose, and methyl-α-glucose in *S. pombe* (Figure S6). Beyond modulating cell growth and proliferation, nutrient availability and cellular metabolic states control the transition from asexual to sexual reproduction in yeasts (Kawamukai 2024). We thus tested the ability of *S. pombe inv2*Δ deletion mutants to undergo mating, meiosis, and sporulation on various media. Microscopic examination of *inv2*Δ mutants revealed a noticeable decrease in the number of zygotes and spore-containing asci, as compared to WT controls (Figure 6A). This phenotype was specifically observed in media containing malt extract, but not upon nitrogen and amino-acid starvation (Figure S7A). Quantification showed that the mating and sporulation defects, although modest, were reproducible across a large number of independent replicates (Figure 6B). To strengthen this observation, we attempted to rescue the observed phenotype by transforming *inv2*Δ mutants with a wild-type copy of the *S. pombe* Inv2 open-reading frame (ORF) fused to YFP and under the control of a tetracycline-inducible promoter, integrated as a single copy plasmid (Lyu et al. 2024). Induction of Inv2-YFP expression fully complemented the mating and sporulation defects (Figure 6C), confirming the specificity of this phenotype and the functionality of the tagged version of Inv2 used in enzymatic assays. In parallel, expression of a catalytically inactive mutant of Inv2, Inv2-D18N, did not rescue mating and sporulation in *inv2*Δ mutants (Figure 6C), indicating that Inv2 enzymatic activity is required for sexual differentiation in *S. pombe*. We next quantified mating and sporulation efficiencies separately. Counting zygotes revealed that the mating step is impaired in *inv2*Δ mutants (Figure 6D). In contrast, analysis of homozygous diploid *inv2*Δ mutants showed that sporulation proceeds with efficiency comparable to that of isogenic WT strains (Figure 6E). Finally, to test whether Inv1 can compensate for the loss of Inv2 function during sexual differentiation, we used a tetracycline-inducible promoter to overexpress a mutant form of Inv1 lacking its signal peptide in an *inv2*Δ background (Figure S7B,C). The resulting defects remained comparable to those observed in *inv2*Δ mutants, indicating that Inv1 cannot substitute for Inv2 during sexual differentiation.

**Figure 6:**
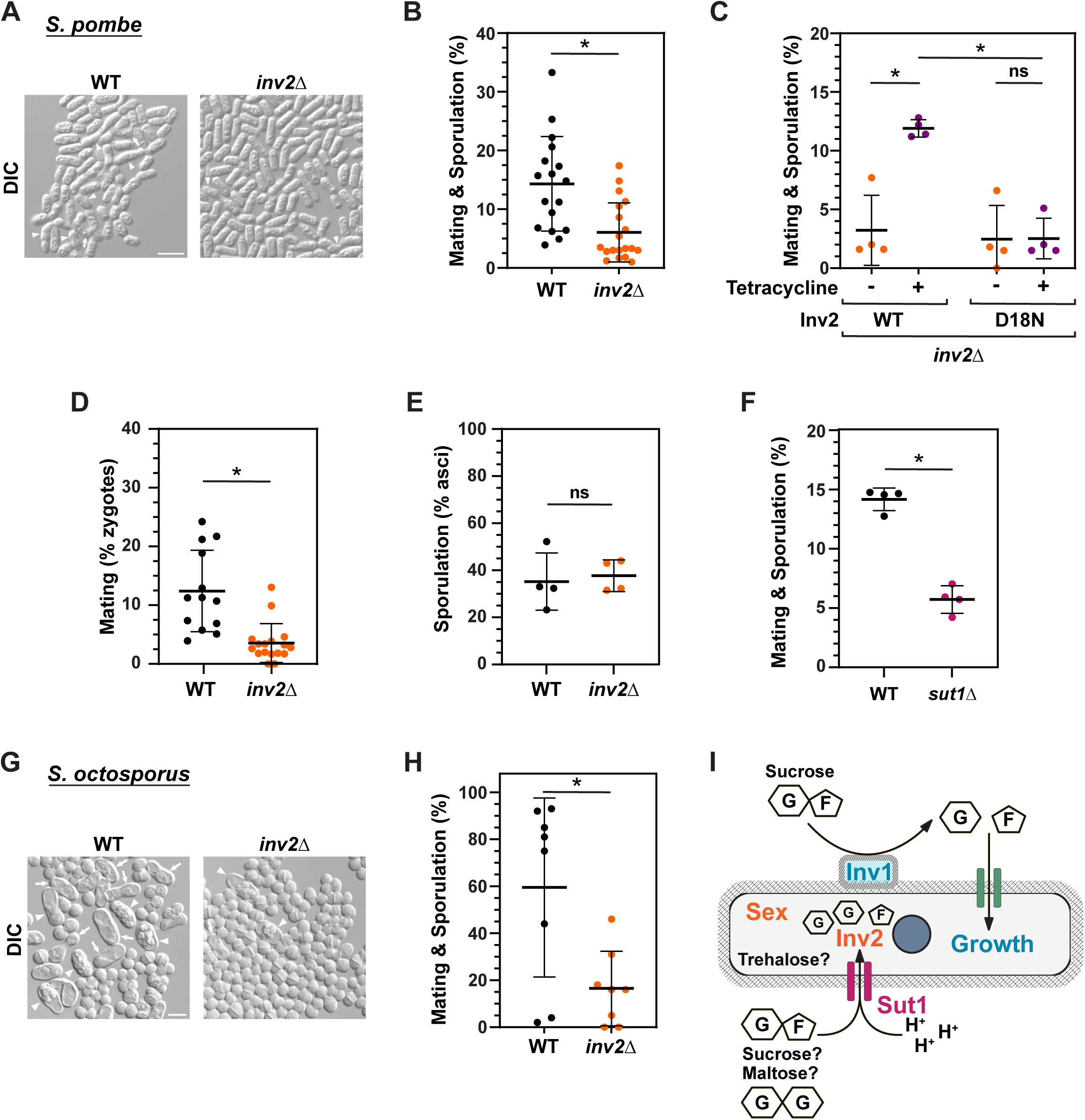
Inv2 is required for sexual reproduction in fission yeasts. (A) Differential interference light microscopy images of *S. pombe inv2*Δ deletion mutants and isogenic WT controls grown on solid malt extract at 28°C for 4 days. (A) White arrows and arrowheads indicate zygotes and asci, respectively. Scale bar: 10 μm. (B) Quantification of the number of zygotes (mating) and asci (sporulation) in WT and *inv2*Δ deletion mutants grown as in (A). Shown are the mean value of at least 17 independent biological replicates, overlaid with individual data points and the SD. Statistical significance was determined by an unpaired, two-tailed t-test. \**p* ≤ 0.05. (C) Mating and sporulation efficiency of *inv2*Δ deletion mutants complemented with either wild-type *inv2* (WT) or an *inv2* catalytic mutant (D18N) ORF (CI) using a tetracycline-inducible promoter. Strains were grown on solid malt extract at 28°C for 4 days before counting. Shown are the mean value of 4 independent biological replicates, overlaid with individual data points and the SD. Statistical significance was determined by two-way ANOVA followed by Tukey’s multiple comparison tests. \**p* ≤ 0.05. (D) Quantification of the number of zygotes to specifically measure mating efficiency in WT and *inv2*Δ deletion mutants grown as in (A). Shown are the mean value of at least 13 independent biological replicates, overlaid with individual data points and the SD. Statistical significance was determined by an unpaired, two-tailed t-test. \**p* ≤ 0.05. (E) Quantification of the number of spore-containing asci to specifically measure meiosis and sporulation efficiency in diploid WT and *inv2*Δ homozygous deletion mutants grown as in (A). Shown are the mean value of 4 independent biological replicates, overlaid with individual data points and the SD. Statistical significance was determined by an unpaired, two-tailed t-test. ns: not significant. (F) Quantification of the number of mating and asci sporulation in WT and *sut1*Δ deletion mutants grown as in (A). Shown are the mean value of 4 independent biological replicates, overlaid with individual data points and the SD. Statistical significance was determined by an unpaired, two-tailed t-test. \**p* ≤ 0.05. (G) Differential interference light microscopy images of *S. octosporus inv2*Δ deletion mutants and isogenic WT controls grown in liquid minimal medium supplemented with malt extract at 30°C for 15 hours. White arrows and arrowheads indicate zygotes and asci, respectively. Scale bar: 10 μm. (H) Quantification of the number of mating and asci sporulation in WT and *inv2*Δ deletion mutants grown as in (G). Shown are the mean value of 8 independent biological replicates, overlaid with individual data points and the SD. Statistical significance was determined by an unpaired, two-tailed t-test. \**p* ≤ 0.05. (I) Invertase gene duplication resulted in either neofunctionalization or subfunctionalization of invertases in the fission yeast clade. See Discussion for details.

The intracellular localization of Inv2 (Figure 4A) prompted us to test whether the putative maltose and sucrose transporter Sut1 also controls sexual reproduction. Similar to *inv2*Δ, *sut1*Δ mutants showed about a two-fold decrease in the number of zygotes and asci in malt extract (Figure 6F), suggesting that uptake of maltose, sucrose, or both disaccharides is important to trigger mating, meiosis and sporulation in *S. pombe*, at least in this metabolic condition. Finally, we found that sexual differentiation was also impaired in *S. octosporus inv2*Δ deletion mutants, as compared to WT isogenic controls (Figure 6G,H). Altogether, our results demonstrate that Inv2 contribute to sexual differentiation in fission yeasts.

## Discussion

Here we identified genomic signatures associated with fission yeast evolution and life history, leveraging recently discovered species and newly generated high-quality genome assemblies. Examining patterns of gene gain and loss, we uncovered a gene duplication event that preceded the lineage-specific loss of an enzyme important for carbon metabolism. One paralog, the Inv1 invertase, functions at the cell wall to hydrolyse sucrose as a carbon source fuelling vegetative growth, whereas the other invertase, Inv2, acts inside the cell to promote sexual differentiation (Figure 6I). Altogether, our comparative analysis of a seemingly well-characterized enzyme reveals unexpected diversity in sucrose utilization strategies among fission yeasts, possibly reflecting their adaptation to habitats with distinct sugar compositions.

### Fate of fission yeast invertase duplication

We accumulated phylogenetic, biochemical, and functional evidence that Inv1 and Inv2 diverge in their enzymatic properties, subcellular localization, expression regulation, and physiological roles (Figure 6I). Inv1 is the canonical yeast invertase essential for growth on sucrose. It displays high affinity for its substrate, high catalytic efficiency with an acidic pH optimum, glycosylation, cell wall localization, and glucose repression, as expected from earlier studies (Mitchison et al. 1969; Moreno et al. 1985; Moreno et al. 1990b; Tanaka et al. 1998; Ahn et al. 2012; Aburigal et al. 2014). In contrast, Inv2 exhibits lower affinity and catalytic efficiency, a higher pH optimum, and lacks glucose regulation, glycosylation, and a signal peptide for secretion. Two studies previously reported the presence of two invertase activities in *S. pombe*, but it remains unclear whether one corresponds to Inv2 because the optimal pH and substrate affinity (*K_m_*) differ substantially from our measurements (Figure 3 and (Mitchison et al. 1969; Moreno et al. 1985)). This discrepancy likely arises from the use of non-purified enzyme preparations, which prevents controlling the substrate-to-enzyme ratio necessary for rigorous kinetic analyses. Consistent with this interpretation, the reported values for the derepressed external invertase activity vary between those studies and also differ from our measurements (Figure 3C,D).

Why is Inv2 catalytically less efficient than Inv1? Structural studies of yeast invertase have shown that it folds into a catalytic β-propeller domain, connected to a C-terminal β-sandwich domain characteristic of GH32 enzymes (Figure S2 and (Álvaro-Benito et al. 2010; Sainz-Polo et al. 2013)). Although the β-sandwich domain is less conserved, it plays a critical role in forming the unusual quaternary structure of a tetramer of dimers. Structure-function analyses further indicate that invertase dimerization enhances catalytic efficiency and substrate recognition through intermolecular polar interactions. Inv2 may, therefore, adopt a distinct oligomeric topology, although the poor conservation of the β-sandwich domain does not allow identification of specific regions or residues that might account for differences in quaternary structure between Inv1 and Inv2 (Figure S2). We propose that the selective pressure for oligomerization, as well as glycosylation and catalytic efficiency, has been stronger for Inv1 than for Inv2, given that Inv1 is secreted. Indeed, although periplasmic localization eliminates the need for sucrose import, it also results in the release of glucose and fructose, which can diffuse away and benefit competing ‘cheaters’ (see Figure 5A and (Gore et al. 2009; Lindsay et al. 2024)). This constraint may have favored the evolution of a more efficient catalytic domain, as well as additional processing steps, such as oligomer formation and extensive glycosylation, to stably anchor the external invertase into the glycoprotein matrix of the cell wall.

Interestingly, plant cytosolic invertases also show lower catalytic efficiency than their extracellular or vacuolar counterparts, likely related to their subcellular localization. Indeed, they differ in their pH optima, with intracellular enzymes classified as neutral or alkaline and secreted or vacuolar forms as acidic. Similarly, Inv2 exhibits an optimum close to neutral pH, matching the intracellular pH of *S. pombe* (Karagiannis and Young 2001), whereas Inv1 is acidic and becomes inactive under such condition (Figure 3B). Therefore, differences in fission yeast invertase catalytic efficiencies may reflect constraints not only imposed by ecological context, but also by differences in physicochemical conditions between subcellular compartments.

Inv1 and Inv2 also differ in their biological roles. Unlike Inv1, Inv2 is dispensable for sucrose assimilation in fission yeasts. Although we cannot exclude the possibility that Inv2 contributes to sucrose utilization under conditions not tested here, adding the Inv1 secretion signal to Inv2 is insufficient to restore growth in the absence of Inv1, suggesting substantial functional divergence. Specifically, the absence of detectable Inv2 secretion suggests that it diverged to the extent that it no longer possesses the sequence features required for endoplasmic reticulum processing, glycosylation, and possibly oligomerization, thereby losing its extracellular growth-promoting function. Instead, our genetic analyses demonstrate a specific role for Inv2 during the mating stage of sexual reproduction in fission yeast. In addition, redirecting Inv1 to an intracellular localization failed to compensate for the loss of Inv2, further suggesting that the two paralogs diverge functionally.

In yeasts, nutrient availability controls the switch from vegetative growth to sexual differentiation. In *S. pombe*, nutrient starvation induces G1 arrest and exit from the cell cycle, followed by conjugation of two haploids cells into a diploid zygote, which immediately undergoes meiosis and differentiation into spores. Remarkably, decades of work established that nitrogen quality and quantity are the primary cues governing this developmental switch in *S. pombe* (Kawamukai 2024). In contrast, the contribution of carbon sources remains comparatively less understood, although previous studies showed that malt extract, glycerol, or galactose can induce sexual differentiation in *S. pombe* even in the presence of nitrogen (Egel 1971; Malecki et al. 2016; Du et al. 2026). Consequently, the mechanistic basis by which Inv2-mediated sucrose hydrolysis influences mating remains speculative.

One possibility is that fission yeasts synthesize sucrose from glucose and fructose, although this appears unlikely because canonical sucrose-synthesizing enzymes are plant-specific and we did not identify clear orthologs in fission yeasts by BLAST-based homology searches. A second possibility is that our sporulation media contains traces of sucrose that would be imported by Sut1 before hydrolysis by Inv2. Indeed, disrupting the putative sucrose transporter Sut1 phenocopies the loss of Inv2. The resulting glucose and fructose produced intracellularly would serve either as metabolic substrates to produce the energy required to complete sexual reproduction or, as demonstrated in plants, as a signaling molecule regulating gene expression and other cellular processes (Ruan 2014). However, supplementing the standard *S. pombe* sporulation medium with sucrose did not trigger mating and sporulation defects in *inv2*Δ mutants (Figure S7A). Likewise, adding glucose and fructose either individually or in combination did not rescue the mating and sporulation defects of *inv2*Δ mutants in *S. pombe* and *S. octosporus* (Figure S7D,E). A third possibility is that Inv2 hydrolyzes a broader range of disaccharides, such as maltose and trehalose, with possible relevance to fission yeast sexual reproduction. First, maltose is the predominant carbon source in malt extract, a component shared by the sporulation media used for *S. pombe* and *S. octosporus* (Figure 6). Supporting this possibility, when expressed heterologously, Sut1 can transport both maltose and sucrose, displaying higher affinity for the former (Reinders and Ward 2001). Second, in *S. pombe*, intracellular trehalose accumulates to high levels under environmental stress, including nitrogen starvation, and its degradation by the neutral trehalase Ntp1 is specifically required for spore germination (Cansado et al. 1998; Beltran et al. 2000; Sajiki et al. 2013; Sakai et al. 2024). However, we found no evidence that Inv2 utilizes either maltose or trehalose to support growth *in vivo* (Figure S6) and the presence of intracellular maltase and trehalase in *S. pombe* makes this interpretation less parsimonious. Beyond disaccharides, Inv2 could potentially hydrolyse fructose-containing oligosaccharides, such as 1-kestose or nystose, as reported for *A. niger* SucB, although no dedicated transporters are known in fission yeasts. In addition, some GH32 enzymes catalyze transfructosylation reactions that generate fructose oligomers (Trollope et al. 2015), but this activity is detectable only at very high sucrose concentrations that are even less likely to occur intracellularly under conditions favouring fission yeast sexual differentiation. Finally, some glycoside hydrolases can degrade plant-derived cyanogenic glycosides, for example during microbiome-mediated detoxification of nectar in bees (Motta et al. 2022), raising the possibility that Inv2 might hydrolyze even more atypical substrates.

In conclusion, we propose that duplication of the fission yeast invertase gene gave rise to a new function that was retained because of its importance for sexual reproduction under specific nutrient-starvation conditions. While Inv1 retained the ancestral role in sucrose-assimilation, Inv2 acquired a role in salvaging low intracellular sucrose levels, either to supply energy or to modulate signalling pathways that influence commitment to sexual reproduction (Figure 6I). Consistent with this functional divergence, Inv1 expression is tightly regulated by glucose levels, which requires multiple regulatory elements spread over large distances (Ahn et al. 2012), whereas Inv2 appears constitutively expressed (Figure 4 and S8). Although directly linking the functions of duplicated genes to ecological adaptation remains challenging, we speculate that this new function of Inv2 emerged as an adaptation to sucrose-poor habitats encountered by the last common ancestor of fission yeasts. Species within the *Sjv* clade have secondarily lost this function (Figure 2A), possibly because they thrived on sucrose-rich substrates such as fruits, as suggested by field surveys (Benito Saez et al. 2013; Brysch-Herzberg et al. 2022; da Silva et al. 2024). Conversely, the canonical extracellular invertase Inv1 was lost in the *Scols* lineage (Figure 2A), consistent with their recurrent isolation from bee food provisions, such as honey and pollen bread, where sucrose levels are low due to enzymatic processing by bees. Finally, the generalist fission species *S. pombe* retained both functions, potentially reflecting adaptation to a broader and more variable ecological landscape.

### Fungal invertase diversity

*S. pombe* represents a rare example of a fungal species that possesses two functional invertases. Indeed, most fungi rely on a single enzyme for sucrose metabolism and studies of *S. cerevisiae* Suc2 served as a paradigm to understand its properties. The canonical yeast invertase is targeted to the cell wall as a heavily glycosylated octamer, is essential for growth on sucrose, and is encoded by a gene under strong glucose-catabolite repression. Early work also detected intracellular invertase activity, later shown to originate from a shorter mRNA isoform generated by alternative transcription start site selection that excludes the N-terminal signal peptide (Perlman and Halvorson 1981; Carlson and Botstein 1982). However, deleting the Suc2 signal peptide abolishes both secretion and sucrose fermentation, demonstrating that the cytosolic isoform is not involved in sucrose utilization (Perlman et al. 1986). Whether it contributes to *S. cerevisiae* fitness in other ways or is purely neutral remains unknown.

Beyond this case, bioinformatic analyses identified putative intracellular invertases across fungal genomes, defined by the absence of a predicted secretion signal within GH32 family enzymes (Figure 2A and (Goosen et al. 2007)). Of these, *Aspergillus niger* SucB is the only cytosolic invertase characterized in detail. SucB displays high catalytic efficiency and strong affinity for sucrose, hydrolyzes longer fructo-oligosaccharides, and can catalyze oligomerization reactions. Surprisingly, although SucB expression is under glucose-catabolite repression, its disruption does not affect growth on sucrose but, rather, affects sporulation timing (Goosen et al. 2007). Thus, similar to *S. pombe* Inv2, SucB regulates sexual reproduction rather than vegetative growth. Unlike SucB, however, Inv2 expression is not regulated by glucose levels, has a pH optimum close to the intracellular pH, and shows substantial lower affinity for sucrose. Interestingly, cytoplasmic invertases are widespread in plants, although they belong to the GH100 family of glucosidases and share no sequence or structural homology with extracellular or vacuolar invertases (Ruan 2014; Wan et al. 2018). Despite this evolutionary divergence, *S. pombe* Inv2 shows notable functional resemblance to plant cytosolic invertases, as both lack glycosylation, exhibit weak catalytic activity, and operate optimally near neutral pH.

Altogether, these observations indicate that functional intracellular invertases exist in fungi, arising through gene duplication followed, most likely, by neofunctionalization. Indeed, secreted invertases that are regulated by glucose levels and essential for sucrose assimilation are thought to be widespread across fungi ((Parrent et al. 2009; Manoochehri et al. 2020) and Figure S1C). Thus, the most parsimonious explanation is that invertase paralogs were independently retained in *S. pombe* and *A. niger* because, in each lineage, the duplicated invertase gene acquired a new function associated with intracellular relocalization, linking the availability of carbon sources to the switch between asexual and sexual reproduction. Nevertheless, we cannot rule out alternative scenarios, such as subfunctionalization (Conant and Wolfe 2008; Panchy et al. 2016), but the limited number of fission yeast species and long phylogenetic distances separating related taxa currently preclude reconstruction of the ancestral function. Interestingly, although these analyses remain predictive and require functional validation, invertases lacking a secretion signal may be more common than previously appreciated ((Parrent et al. 2009) and Figure 2A). Finally, the presence of two *S. cerevisiae* invertase isoforms with distinct subcellular localizations provides a striking example of how a single gene can generate both secreted and intracellular activities, although, to our knowledge, the function of the intracellular isoform remains unknown.

### Genomic signatures of fission yeast life history

Knowledge of the evolutionary trajectory and ecology of model organisms is important for understanding fundamental biological processes. Unlike other model species, relatively little is known about the natural history of *S. pombe* and the broader fission yeast clade. Their sporadic occurrence in natural microbial communities suggests that these species are adapted to specific ecological niches, yet the underlying drivers of this specialization remain poorly understood. The recent discovery of additional fission yeast species, together with the availability of high-quality genomes assemblies, now provides an unprecedented opportunity to identify the genetic basis of these evolutionary adaptations.

Comparative ontology of gene gain and loss events revealed that metabolic genes exhibit the largest interspecific variation within this clade, supporting the hypothesis that nutrient availability and assimilation are major selective pressures shaping their genomes. Notably, the profound metabolic rewiring observed in *S. japonicus* is consistent with its loss of respiration and may have been driven by stable, low-complexity nutrient environments. Likewise, both the *Sjv* and *Scols* lineages show substantial metabolic changes, suggesting possible ecological specialization, for example through occupation of niches with partially overlapping abiotic and biotic parameters. Similarly, the loss of RNA-metabolism genes in these lineages may reflect distinct strategies for regulating gene expression and the haploid-diploid life cycle in response to nutrient availability. In contrast, *S. pombe* retained a broader metabolic repertoire, consistent with its ability to thrive across diverse habitats. Furthermore, this species seemingly expanded its metabolic capacities, notably through HGT, possibly to facilitate the assimilation of a broader range of substrates and to adapt to fluctuating environments. Taken together, the lineage-specific gene gains and losses identified here represent a valuable starting point for future functional studies aimed at dissecting the genetic mechanisms underlying ecological adaptation in fission yeasts.

## Materials & Methods

### Genome analysis

Two studies examined the orthology relationships between *Schizosaccharomyces* species: *S. japonicus* (4198 protein-coding genes, *Sj*), *S. pombe* (5123 protein-coding genes, *Sp*), *S. cryophilus* (5056 protein-coding genes, *Sc*), *S. octosporus* (5069 protein-coding genes, *So*) and *S. osmophilus* (5098 protein-coding genes, *Ss*) (Rhind et al. 2011; Jia et al. 2023). We revisited these relationships by including genomes of two recently discovered species: *S. versatilis* (*Sv*) (Brysch-Herzberg et al. 2024; Etherington et al. 2024) and *S. lindneri* (*Sl*) (Brysch-Herzberg et al. 2023), which are closely related to *Sj* and *So*, respectively. Taphrinomycotina genome and protein sequences were retrieved from NCBI (www.ncbi.nlm.nih.gov) and used as local databases for the BLAST search tools available in the Geneious 11.1.5 software package (Biomatters, www.geneious.com/). We identified orthologous proteins by reciprocal BLASTP searches using the query-centric alignment method and selecting the single best-scoring hit in each species (Table S1). In most cases, the best hits corresponded to unique genes whose scores were fully consistent with the phylogeny of the seven fission yeast species. In instances where gene duplications were detected, orthology relationships were further examined using synteny analyses and maximum-likelihood phylogenetic analyses of multiple sequence alignments.

For genes missing from one or several species, we used TBLASTN searches to check which genes are present in genomes but not annotated. We identified 8 unannotated genes in *Sj*, 134 in *Sv*, 13 in *Sc*, 250 in *Sl*, 26 in *So* and 6 in *Ss*. We also searched for hits in more distant species by BLASTP searches in NCBI ClusteredNR databases restricted either to Taphrinomycotina (taxid:451866), Saccharomyceta (taxid:716545), Fungi incertæ cedis (taxid:112252) or Bacteria (taxid 2). These analyses allowed inferring the presence or absence of a homologous gene in distant clades, and thus discriminating between a loss or a gain event in fission yeast species (Table S2). The presence of a homolog in anciently diverged species indicated that the gene was likely lost from fission yeasts. Conversely, genes without any distant homolog were classified as fission yeast gain events. We considered each loss and gain event as unique independently of duplications that may have occurred in each species. Genes considered as “dubious” in PomBase (n=65) (Carme et al. 2026) and JaponicusDB (n=41) (Rutherford et al. 2022) were excluded from the final counts. Finally, TBLASTN searches also allowed identifying horizontal gene transfer (HGT) events from Bacteria, by comparing similarities of best hits in either taxon (Table S1).

For ontology analyses, we extracted GO annotations from PomBase (Table S2) and used GO Slim BP categories to categorize genes gained or lost in *Sjv*, *Spcols*, *Sp* or *Scols*. All events were captured with 40 different BP, which we grouped into 10 broader categories (Table S2), before counting gains and losses for each GO BP category and species group. Heatmaps were constructed with Morpheus (software.broadinstitute.org/morpheus/), using column hierarchical clustering, with Euclidian distance and average linkage as settings.

### Phylogenetic Analyses

Amino acid sequences were aligned using MAFFT v7.45, available in the Geneious 11.1.5 package (Katoh and Standley 2013). Multiple Sequence Alignments (MSAs) were processed by Block Mapping and Processing by Entropy (BMGE) v1.12.1 with a 0.7 cut-off value (Criscuolo and Gribaldo 2010). Phylogenetic trees were estimated by the maximum-likelihood (PhyML; https://ngphylogeny.fr, (Lemoine et al. 2019)) and MrBayes (Geneious 11.1.5). PhyML v3.3_1 was set up using estimated proportion of invariant sites and gamma model (4 categories) parameters. ML trees were optimized for topology, length and rate and were generated using the best of nearest-neighbour interchange and subtree-pruning-regrafting tree search algorithms, with 100 bootstrap replicates. MrBayes consensus trees were generated after two independent runs of four Markov chains for 1,100,000 generations sampled every 200 generations, with sampled trees from the first 100,000 generations discarded as “burn-in”. Newick trees were visualized and exported with FigTree v1.4.4 (https://github.com/rambaut/figtree/releases).

### Fission yeast procedures

Standard rich (YES) and minimal (EMM) media (Forsburg and Rhind 2006) were used to propagate *S. pombe* and *S. japonicus* strains. *S. octosporus* strains were maintained in YES medium supplemented with 30% glucose (w/v). To test the effect of media acidification on sucrose assimilation, the pH of EMM was lowered from 5.5 to 3.5 using saturated HCl. To inhibit respiration, cells were treated with 5 μg/mL antimycin A (AntA, A8674, Sigma). To prevent fermentation and stimulate respiratory growth, cells were grown in EMM containing 0.08% glucose (w/v), as described in (Takeda et al. 2015). Hyperosmolarity was tested by growing cells in EMM containing 10-40% glucose or sucrose. For Inv1 induction, cells were inoculated in high-glucose medium (YES or EMM supplemented with 3% glucose) and grown at 32°C to mid-log phase (∼0.5 x 10^7^ cells/ml). Cells were then harvested by vacuum-based filtration, washed twice using 1X PBS at room temperature, resuspended in either high- or low-glucose media (YES or EMM with 0.1% glucose and 3% glycerol), and incubated at 32°C for the indicated time points. Induction of the *enotet* promoter (Lyu et al. 2024) was performed by adding 2.5 μg/mL anhydro-tetracycline (10792081 Thermo Scientific) resuspended in DMSO to liquid rich media for growth assays or solid malt extract for mating assays.

For all species, qualitative fitness assays were performed by inoculating single colonies into liquid media and dropping either 10-fold serial dilutions on solid media or a 35-fold dilution in ID32C strips (API medium, Biomerieux). For *S. pombe*, quantitative fitness assays were performed by monitoring growth using a microplate absorbance reader (FLUOstar Omega, BMG Labtech or Epoch2, Agilent BioTek), with the following parameters: 32°C temperature, 100 rpm double-orbital shaking, 6 mm spiral averaging, and 600 nm top-optic measurement. Quantitative parameters were obtained by fitting a reparametrized Gompertz non-linear regression model to each growth curve using GraphPad Prism, as described in (Lo Grasso et al. 2023):

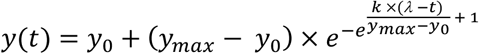

where *y*(*t*) represents the natural logarithm of the optical density (OD) of the culture, a proxy of yeast biomass; *y*_0_ and *y_max_* are the starting and maximum biomass (*OD*_600_), respectively; *k* the maximal specific growth rate (ℎ^−1^); *λ* the lag phase duration (ℎ); and *t* the time (ℎ).

For *S. pombe*, mating assays were performed by resuspending yeast *h^90^*, *h^+^*, and *h^-^*cells in 10 μl sterilized water before spotting onto solid bacteriological-grade malt extract (ME, Gibco, 218630). For *S. octosporus*, cells were spotted on a 1:1 mixture of malt extract and yeast morphology agar (MYA, Formedium CYN0102). After incubation at 28°C for 4 days for *S. pombe* and for 2 days for *S. octosporus*, the number of vegetative cells (V), zygotes (Z), asci (A), and free spores (S) were counted using a differential interference contrast light microscope. The efficiency of sexual differentiation (%) was calculated as follows: (2Z + 2A + S/2) / (V + 2Z + 2A + S/2) x 100 for *S. pombe*; (2Z + 2A + S/4) / (V + 2Z + 2A + S/4) x 100 for *S. octosporus*. At least 100 cells were counted for each independent biological replicate.

### Strain construction

All strains used in this study are listed in Table S4. *S. pombe* strains were constructed by standard procedures, using either chemical transformation or genetic crosses (Forsburg and Rhind 2006). Genetic crosses were performed by mating strains at 28°C on SPAS medium. Diploid strains were obtained from crosses on SPAS medium for 16 hours at 28°C, before selection through purification on minimal media lacking adenine (EMMc-Ade) and mating assays on solid malt extract media. Strains with gene deletions were constructed by PCR-based gene targeting of the respective open reading frame (ORF) with *kanMX6*, *natMX6*, or *hphMX6* cassettes, amplified from pFA6a backbone plasmids (Bahler et al. 1998; Hentges et al. 2005). PCR amplification was performed using primers of 100 bases, with 80 bases to direct homologous recombination. For C-terminal epitope tagging of Inv1 and Inv2, CRISPR-Cas9-mediated genome editing was used, as described in (Zhang et al. 2018). DNA fragments used for homologous recombination were generated by PCR. Primers were designed using the online fission yeast database, PomBase (Rutherford et al. 2024), and are listed in Table S6.

For complementation experiments, the *inv2* ORF was amplified from genomic DNA in two steps using DHO3417, 3431, 3432, and 3426 and subcloned into NheI-digested DHB317 (pDB5319, (Lyu et al. 2024)) by Gibson assembly (E2621L, New England Biolabs) to generate DHB369. In this plasmid, the full-length *S. pombe inv2* ORF is fused to the YFP variant mECitrine at its C-terminal end and placed under the control of the *enotet* tetracycline-inducible promoter. DHB369 was used as a template for quick-change site-directed mutagenesis to create a G-to-A mutation at position +52 (+1 defined as the A from the initiating codon) to replace Asp18 with an Asn18 residue, using DHO3768 and 3769, creating DHB418. This mutation abolishes *S. cerevisiae* Suc2 catalytic activity *in vitro* (Reddy and Maley 1990). We generated DHB370 using a DNA fragment corresponding to the signal peptide of *S. pombe* Inv1 (+1 to +79) at the 5’ end of the *inv2* ORF, amplified using DHO3418 and 3427, assembled with the *inv2* ORF amplified using DHO3417, 3431, 3432, and 3433 into NheI-digested DHB317. The *inv1* ORF was amplified from genomic DNA using primers designed to remove its N-terminal signal peptide sequence, DHO3700 and 3701, and subcloned into pUC19 digested with HindIII and EcoRI to generate DHB416. Then *inv1* was re-amplified using DHO3999 and 4000 and subcloned into NheI-digested DHB317 to generate DHB442, in which the *S. pombe* Inv1 ORF lacking the signal peptide (+80 to +581) is fused to the YFP variant mECitrine at its C-terminal end and placed under the control of a tetracycline-inducible promoter. All plasmids were digested with NotI before integration 3’ of the *ade6* 3’ UTR in *S. pombe* using lithium-acetate transformation. All plasmids and primer sequences are listed in Table S5 and S6, respectively.

*S. japonicus* and *S. octosporus* gene editing was performed by stable integration of plasmids into the genome. DNA fragments used for homologous recombination were obtained by gene synthesis (Integrated DNA Technologies) and subcloned by Gibson assembly (E2621L, New England Biolabs). All fragments and PCR primers were designed using the online yeast database EnsemblFungi (Yates et al. 2022). We first obtained DHB283 by inserting a multiple cloning site into a modified pRS316 backbone (Addgene #74081, (Al-Sady et al. 2016)), upstream of the superfolder GFP ORF (sfGFP), the *S. pombe ura4* terminator, and the *hphMX6* cassette. Then, for *S. japonicus inv1* gene deletion, a DNA fragment comprising 300-bp of 5’ and 3’ homologous sequences was designed to precisely delete its ORF (from the ATG to the stop codon) and assembled into the XbaI site to generate DHB285. For *S. octosporus inv2* gene deletion, DNA fragments comprising 1000-bp of 5’ and 3’ homologous sequences were designed in two steps to precisely delete its ORF (from the ATG to the stop codon), amplified by PCR using DHO2790, 2791, 3086, and 3087, and assembled into the XbaI site to generate DHB287. The 5’-to-3’ order of homologous sequences was inverted within the final plasmids to obtain single-copy recombination into the genome (Vještica et al. 2020), upon linearization with StuI and yeast transformation. Plasmids were verified by Sanger and Oxford nanopore sequencing and transformed by electroporation using previously published protocols (Aoki et al. 2010; Seike and Niki 2017). For *S. octosporus*, 50 mL of cells were grown in YE with 30% glucose at 30°C to mid-log phase (∼3 x 10^7^ cells/mL), harvested by centrifugation (3,500 rpm, 3 mins, 4°C), and washed thrice with ice-cold sterilized water. Cells were resuspended in 5 mL of 1M sorbitol and 125 μL of 1M dithiothreitol (DTT), incubated for 15 mins at 30°C without shaking, spun down at 3,500 rpm for 3 mins at 4°C, washed in 1 mL of 1M sorbitol, and resuspended in 100 μL of 1M sorbitol, to which up to 5 μg of transforming DNA was added, mixed with 2 μg of sonicated salmon sperm DNA. After incubation on ice for 5 mins, the mixture was electroporated using a MicroPulser electroporator (2 mm cuvette, 3.0 kV, Bio-Rad), and then resuspended in 1 mL 1M sorbitol and 8 mL of YE with 30% glucose, before incubation for at least 16 hours at 30°C with shaking and plating onto selective agar medium. Transformants were screened for correct integration by colony-PCR and, when appropriate, verified by Western blotting. For each transformation, 2-4 individual clones were purified and analyzed. All plasmids and primer sequences are listed in Table S5 and S6, respectively.

### Live fluorescence microscopy

About 10 mL of *inv1-sfGFP* and *inv2-sfGFP* EMM cultures or 10 mL of *inv1*Δ *tetON-SP-inv1-inv2mECitrine* YES cultures with and without tetracycline were harvested by centrifugation. Cells were resuspended in 1 mL 1X PBS containing 1.5 µg/mL DAPI (D9542, Sigma) before a 30 min incubation at room temperature in the dark and image acquisition either on a Zeiss LSM 980 NLO confocal microscope (Figure 4A) or a Zeiss AxioImager Z2 upright epifluorescence microscope (Figure S5B).

### Fluorescence measurements

Total fluorescent levels of tetracycline-inducible Inv1-YFP and Inv2-sfGFP were measured using a microplate absorbance reader (FLUOstar Omega, BMG Labtech or Epoch2, Agilent BioTek) from solid media drops of yeast cells resuspended in 200 μL water. Fluorescence levels were normalized to background fluorescence in untagged strains and total yeast cell concentration by measuring absorbance (OD600).

### RT-qPCR analysis

Reverse transcription and quantitative PCR analyses of cDNA were performed using RNA extracted from 50 mL of exponentially growing cells, as described in (Laboucarié et al. 2017), and according to the MIQE guidelines (Bustin et al. 2009). Briefly, total RNA was purified using hot, acidic phenol and contaminating DNA was removed by DNase I digestion, using the TURBO DNA-free™ kit (AM1907, Ambion). 1 μg of RNA was then reverse transcribed (RT) at 55°C with random hexanucleotide primers, using the SuperScript III First-Strand System (18080051, ThermoFisher Scientific). Fluorescence-based quantitative PCR was performed with SYBR Green on a CFX Opus 384 Real-Time PCR System (Bio-Rad) and used to calculate relative cDNA quantities, from the slope produced by standard curves for each primer pair, in each experiment. DNase-treated RNA samples were used as controls for the presence of genomic DNA contaminants. Standard curve slopes were comprised between -3.5 (90% efficiency) and -3.15 (110% efficiency), with an *r*^2^ > 0.9. All primer sequences are listed in Table S6.

### Protein extraction

Protein extracts were prepared as described in (Laboucarié et al. 2017). Briefly, 25 mL cultures of exponentially growing cells were homogenized by glass bead-beating in a FastPrep (MP Biomedicals). Proteins extracted using either standard lysis buffer (WEB: 40 mM HEPES-NaOH pH 7.4, 350 mM NaCl, 0.1% NP40, and 10% glycerol), supplemented with protease inhibitors, including cOmplete EDTA-free cocktails tablets (04693132001, Roche), 1 mM PMSF (P7626, Sigma), 1 µg/ml bestatin (B8385, Sigma), and 1 µg/ml pepstatin A (P5318, Sigma).

### Immunoprecipitation

Whole protein extracts were prepared from 100 mL YES cultures of *inv1-sfGFP* and *inv2-sfGFP* strains in immunoprecipitation (IP) lysis buffer (50 mM HEPES-NaOH pH 7.4, 150 mM NaCl, 0.5% NP40, 1 mM EDTA and 10% glycerol). About 15-20 mg of protein extracts were incubated with 25 μL of GFP-Trap Agarose beads (gta-20, Chromotek) for 2 hours at 4°C. Beads were washed 3 times with IP lysis buffer and eluted with 0.1 M Glycine pH 2.5. Eluates were supplemented with 1 μL of 1 M Tris-base to neutralize the eluted samples to pH 7. Total protein content of the eluates was visualized by SDS-PAGE followed by Coomassie blue staining.

### Western blotting and antibodies

Western blotting was performed using an anti-GFP antibody (TP401, CliniScience). Protein concentrations were measured by the Bradford method and used to load equal amounts of proteins across samples. Ponceau red staining was used to normalize for total protein levels across samples. Acquisition of the chemiluminescence signal was performed using an Odyssey Fc Imager and analyzed with Image Studio™ Lite 4.0 (LI-COR Biosciences).

### Glycosylation assay

Anti-sfGFP immunoprecipitations were denaturated at 100°C during 10 minutes, incubated with 500 Units of Endoglycosidase H (P0702S, New England Biolabs) for 1 hour at 37°C before SDS-PAGE and Western blotting.

### Invertase enzymatic assay

Eluates from anti-sfGFP immunoprecipitations were mixed with sucrose and 0.5 M potassium acetate buffer at pH 4.5 for Inv1 and pH 6.0 for Inv2 in a 100 μL reaction. The mixture was incubated at 50°C during 10 mins for Inv1 and 60 mins for Inv2. To test the linearity of the reaction, Inv2 enzymatic assays were performed over a time course of 90 mins. To test the effect of pH on Inv1 and Inv2 activities, enzymatic assays were performed using potassium acetate buffer adjusted to pH values ranging from 2 to 8. Kinetic assays were performed using increasing concentrations of sucrose, ranging from 0-5.2% (w/v) for Inv1 and 0-50% for Inv2.

Since glucose is the only reducing sugar produced upon sucrose hydrolysis by invertase, glucose concentration was determined using the dinitrosalicylic acid reagent (DNS) and colorimetric measurements, as described in (Wood et al. 2012). DNS was prepared by mixing 10 g/L 3,5-dinitrosalicylic acid (D0550, Sigma), 30 g/L sodium potassium tartrate (217255, Sigma), 16 g/L NaOH. Each enzymatic reaction was mixed with DNS to a final volume of 180 μL, at a 1:20 and 1:10 ratio for Inv1 and Inv2, respectively. The resulting mixture was heated at 100°C for 1 min, cooled down to 20°C for 2 min using a thermocycler, and transferred to a 96-well microplate for measuring absorbance at 540 and 580 nm at 25°C using a spectrophotometer (FLUOstar Omega, BMG Labtech). In each experiment, absolute glucose concentrations were calculated using standard curves, which were obtained by measuring D-glucose solutions of known concentrations at both wavelengths and fitting a linear equation. Quantitative enzymatic parameters were obtained by fitting a Michealis-Menten equation to kinetic curves using GraphPad Prism:

### Quantification and statistical analysis

All statistical tests were performed using GraphPad Prism (version 10). *t*-tests were used when comparing two means, using Welch’s correction when variances were not equal. Two-way analyses of variance (ANOVA) were performed for comparing more than two means, across two distinct variables (for example “genotype” and “treatment”), followed by Tukey *post hoc* pairwise comparisons. A significance level (α) of 0.05 was used *a priori* for all statistical tests. Statistical details of experiments can be found in the figure legends, including the statistical tests used, the minimum value of biological replicates n shown (n = isogenic clones of each strain). Quantitative values are typically represented as individual values (n) overlaid with the mean (black bar) and standard deviation (SD).

## Acknowledgments

We thank Dune Noly and Karim Majzoub for critical reading of the manuscript. We are also grateful to Li-Lin Du for sharing valuable knowledge prior to publication. We thank all members of the Helmlinger laboratory for helpful suggestions and discussions. We thank Kevin Terretaz and the Montpellier Imaging Resource (MRI) for assistance with confocal microscopy. We thank the joint IGMM-CRBM "Yeast Media and Technologies" service for providing us with ready-to-use yeast media. We are indebted to Snezhana Oliferenko for sharing strains and the National Bio-Resource Project (NBRP) – Yeast, Japan for providing the pDB5319 plasmid. A.N. was a recipient of fellowships from the French Ministry for Research and Higher Education (MESR) and the Fondation pour la Recherche Médicale. This work was supported by funds from the CNRS MITI 80Prime to D.H..

## Data Availability Statement

All data supporting the findings of this study are included in the article and its supplementary materials. Strains, plasmids, and source data are available from the corresponding author upon reasonable request. Requests for access to these materials and data will be addressed by the corresponding author.

## Figure Legends

**Supplemental Figure S1: Evolutionary history of fungal invertases.**

(A) Heat map showing percent identity between Schizosaccharomycetes Inv1 and Inv2 amino-acid sequences, compared to *S. cerevisiae* Suc2. Sequences were aligned using MAFFT with the BLOSUM62 scoring matrix. Percent identity values are color-coded according to the scale shown on the right.

(B) Bar plot of average percent identity between Schizosaccharomycetes Inv1, Inv2 and *S. cerevisiae* Suc2. Bars represent mean pairwise identity values overlaid with standard deviations.

(C) Phylogenetic tree of invertases from fungi, bacteria and plants, based on 117 sequences from 109 species selected as best hits from BLASTp searches using *S. pombe* Inv1 and Inv2 as queries. All sequence accessions are listed in Table S3. Sequences were aligned with MAFFT and filtered with BMGE (entropy cutoff = 0.7). The tree was inferred using maximum-likelihood analysis with bootstrap support. Red and blue circles indicate nodes supported by bootstrap values ≥90 and ≥70, respectively. Sequences annotated or predicted to contain a signal peptide (InterPro; https://www.ebi.ac.uk/interpro/ and SignalP 6.0; https://services.healthtech.dtu.dk/services/SignalP-6.0/) are marked with a star. The inset shows fungal taxonomy classes color-coded accordingly. Most fungal invertases segregate into two well-supported groups, designated Cluster I and Cluster II.

**Supplemental Figure S2: Multiple alignments of fission yeast invertases.**

Multiple alignment of *S. pombe* Inv1 (SPCC191.11), *S. pombe* Inv2 (SPAC8E11.01c), *S. japonicus* Inv1 (SJAG_04301), *S. octosporus* Inv2 (SOCG_04818), and *S. cerevisiae* Suc2 (YIL162W) using Clustal Omega (EMBL-EBI) and Jalview 2.11.4.1 for visualization.

Conserved residues are highlighted in shades of blue depending on the percentage of similarity to a consensus sequence.

**Supplemental Figure S3: *S. pombe* Inv2 enzymology controls.**

(A) Direct comparison of invertase activity in affinity purified Inv1-sfGFP and Inv2-sfGFP eluates, along with an eluate from an immunoprecipitation of ‘no-sfGFP’ control strain. Enzymatic velocity values (mM/min) are expressed as a percentage of the maximal measurement and independent experiments are shown as individual data points overlaid with the mean.

(B) Enzyme progress curve of Inv2-sfGFP. An eluate of affinity purified Inv2-sfGFP was incubated for various time points, as indicated, and before measuring glucose concentrations using the DNS reagent. A linear regression fit shows that glucose concentration increases linearly with time (*R^2^*= 0.9799, *P* < 0.0001), demonstrating that enzymatic assays performed during 60 mins for Inv2 correspond to the initial rate phase of the reaction.

**Figure S4: Inv2 fails to compensate for the loss of Inv1.**

(A) Sucrose assimilation of *S. pombe inv1*Δ deletion mutants complemented with Inv2 harbouring the N-terminal signal peptide of Inv1 and placed under the control of a tetracycline-inducible promoter (SP^inv1^-Inv2). Saturated cultures were diluted into liquid minimal media supplemented with 2% glucose or sucrose, and incubated at 32°C for 24 hours. Biomass was measured as a proxy of growth and monitored continuously by measuring absorbance (OD600) at 10 mins intervals. Shown are the mean value of 4 independent biological replicates overlaid with the SD.

(B) Levels and localization of SP^inv1^-Inv2-YFP upon tetracycline induction in strains grown as in (A). fused to the signal peptide of *S. pombe* Inv1. Upper panels show live fluorescence microscopy of SP^inv1^-Inv2-YFP in absence (left) and presence (right) of tetracycline. Bottom panels show differential interference light microscopy images of the corresponding fields. Scale bar: 12 μm.

(C) Ten-fold serial dilutions of exponentially growing cells of the indicated genotypes were spotted in rich medium containing 3% glucose or sucrose and incubated at 32°C for 3 days. White dashed lines represent agar cut out from the plates to prevent diffusion of nutrients.

**Figure S5: Sut1 is not required for sucrose assimilation in *S. pombe*.**

(A,B) Saturated cultures were diluted into liquid minimal media supplemented with 2% glucose or sucrose and incubated at 32°C for 36 hours. Biomass was measured as a proxy of growth and monitored continuously by measuring absorbance (OD600) at 10 mins intervals. For quantitative comparisons between genotypes and conditions, growth rate values (*k*) were computed by fitting a modified logistic growth model to proliferation curves. Shown are the mean value of 4 independent biological replicates overlaid with the SD.

**Figure S6: Inv2 is not required for assimilation of alternative carbon sources in *S. pombe*.**

(A) Ten-fold serial dilutions of exponentially growing cells of the indicated genotypes were spotted in rich medium containing 3% glucose, maltose, trehalose, and methyl α-D-glucose, and incubated at 32°C for up to 6 days. Shown are images that are representative of three independent experiments. White dashed lines in represent agar cut out from the plates to prevent diffusion of disaccharide breakdown products.

(B) Summary of growth of *S. pombe* strains of the indicated genotypes on raffinose, as well as glucose and sucrose as controls, using a miniaturized assimilation test for yeast identification.

**Figure S7: Factors influencing Inv2 function in sexual differentiation.**

(A) Quantification of the number of zygotes (mating) and asci (sporulation) in WT and *inv2*Δ deletion mutants grown on solid SPAS medium supplemented with either 1% glucose and 1% sucrose at 28°C for 2 days. Shown are the mean value of at 6 independent biological replicates, overlaid with individual data points and the SD. Statistical significance was determined by two-way ANOVA followed by Tukey’s multiple comparison tests. ns: not significant.

(B) Mating and sporulation efficiency of *inv2*Δ mutants complemented with a mutant form of Inv1 lacking its N-terminal signal peptide to force intracellular localization (Inv1^woSP^-YFP) using a tetracycline-inducible promoter, along with isogenic WT controls and *inv2*Δ mutants. Strains were grown on solid malt extract at 28°C for 4 days before counting. Shown are the mean value of 3 independent biological replicates, overlaid with individual data points and the SD. Statistical significance was determined by two-way ANOVA followed by Tukey’s multiple comparison tests. \**p* ≤ 0.05; ns: not significant.

(C) Total fluorescence levels of Inv1^woSP^-YFP upon tetracycline induction in strains grown as in (B). Statistical significance was determined by an unpaired, two-tailed t-test. \**p* ≤ 0.05.

(D) Quantification of the number of zygotes (mating) and asci (sporulation) in *S. pombe* WT and *inv2*Δ deletion mutants grown on solid malt extract supplemented with either 1% glucose, 1% fructose, or a mixture of 0.5% glucose and 0.5% fructose at 28°C for 4 days. Shown are the mean value of at least 3 independent biological replicates, overlaid with individual data points and the SD. Statistical significance was determined by two-way ANOVA followed by Tukey’s multiple comparison tests. \**p* ≤ 0.05; ns: not significant.

(E) Quantification of the number of zygotes (mating) and asci (sporulation) in *S. octosporus* WT and *inv2*Δ deletion mutants grown on liquid minimal medium supplemented with malt extract and either 1% glucose or a mixture of 0.5% glucose and 0.5% fructose at 30°C for 2 days. Shown are the mean value of at least 3 independent biological replicates, overlaid with individual data points and the SD. Statistical significance was determined by two-way ANOVA followed by Tukey’s multiple comparison tests. \**p* ≤ 0.05; ns: not significant.

**Figure S8: Inv2 expression levels upon sexual differentiation.**

(A) Quantitative RT-PCR analysis of *inv2* mRNA levels in total RNA extracts from *S. pombe* grown either in rich (solid YES) or starved conditions (solid malt extract) at 28°C for 4 days. *rps1101* served as a control for normalization across samples. Shown are the mean value of 3-4 independent biological replicates, overlaid with individual data points and the SD. Statistical significance was determined by unpaired, two-tailed t-tests. \**p* ≤ 0.05.

(D) Total fluorescence levels of Inv2-sfGFP grown as in (A). Statistical significance was determined by an unpaired, two-tailed t-test. ns: not significant.

