## Supplementary Figures for "Evolutionary diversification of invertase paralogs couples carbon metabolism and sexual reproduction in fission yeasts"

Noly A. *et al.*, Supplemental Figure 1

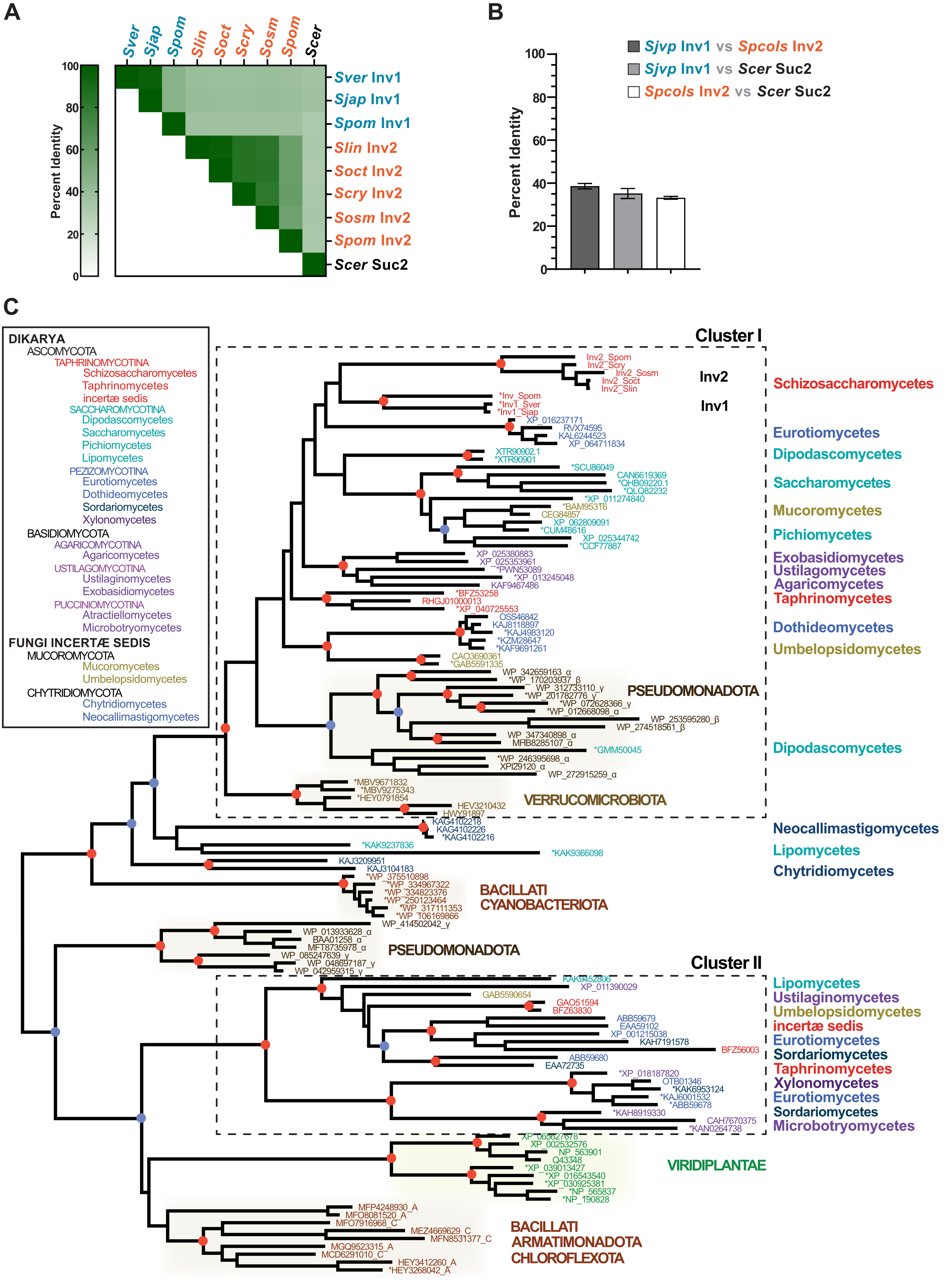

**Noly A. *et al.*, Supplemental Figure 2**

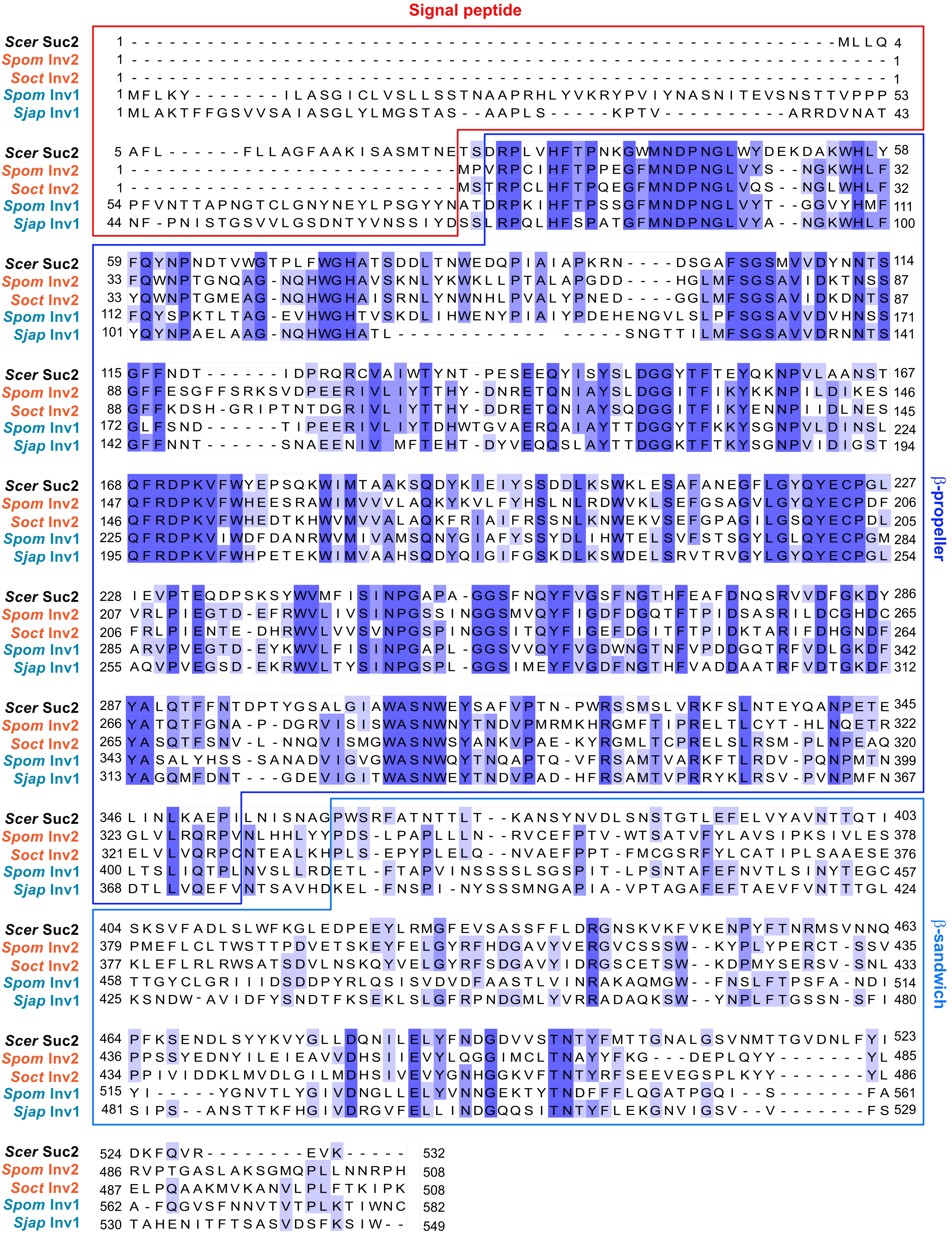

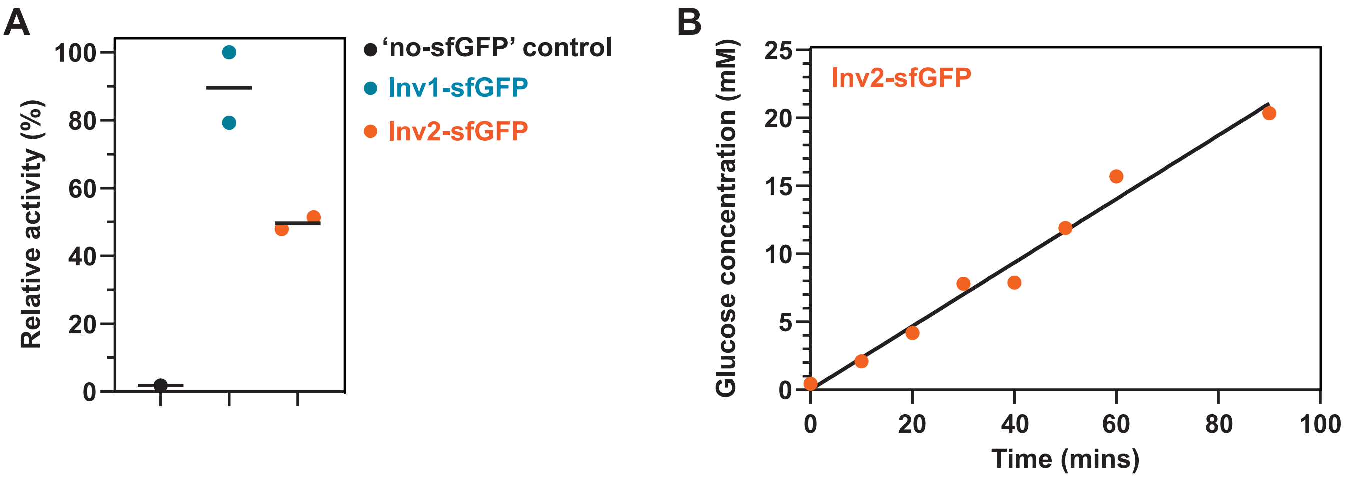
A
Relative activity (%)
100
80
60
40
20
0
'no-sfGFP' control
Inv1-sfGFP
Inv2-sfGFP
B
Glucose concentration (mM)
25
20
15
10
5
0
Inv2-sfGFP
0
20
40
60
80
100
Time (mins)

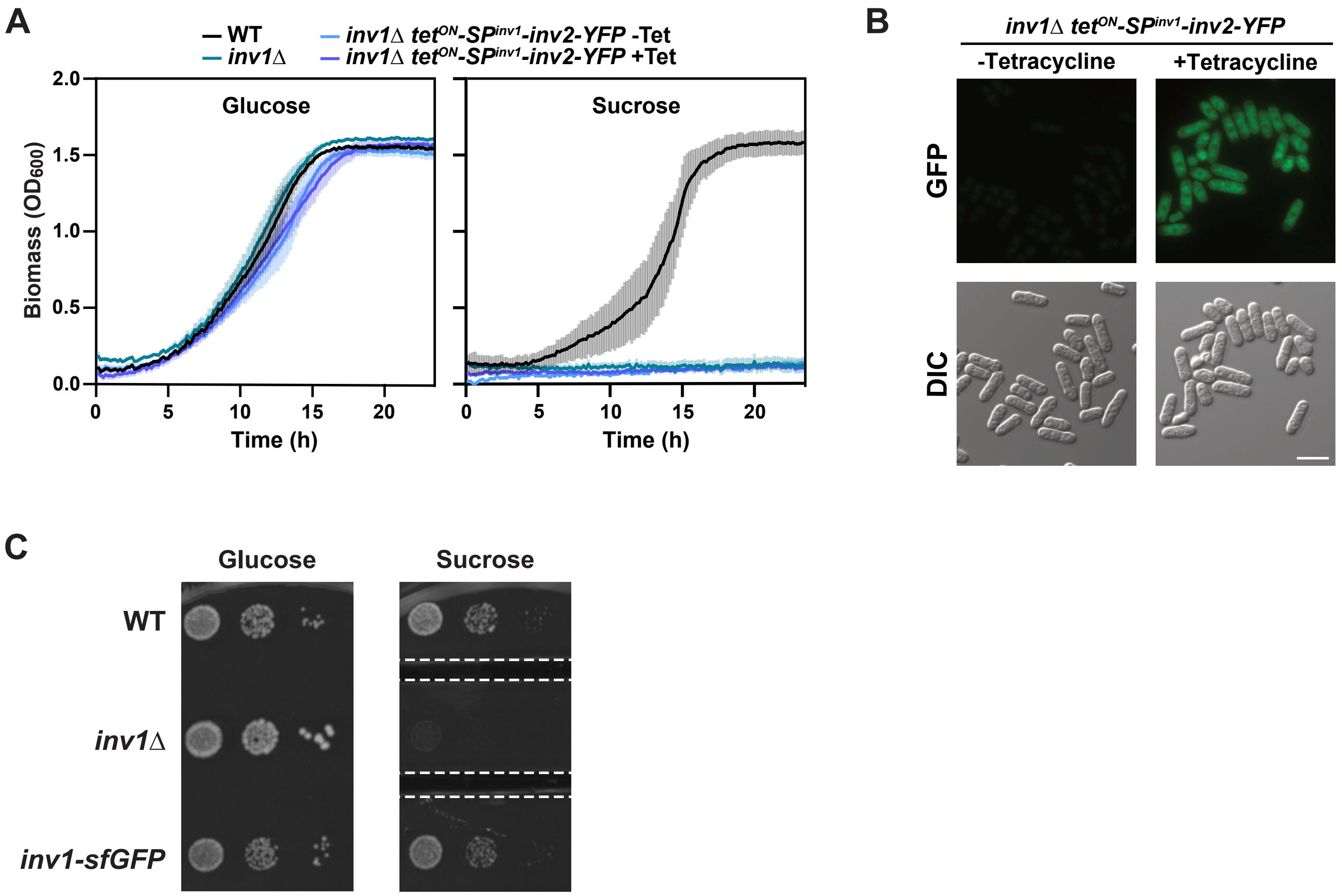
A
WT
inv1Δ
inv1Δ tetON-SPinv1-inv2-YFP -Tet
inv1Δ tetON-SPinv1-inv2-YFP +Tet
Glucose
Sucrose
Biomass (OD600)
2.0
1.5
1.0
0.5
0.0
0
5
10
15
20
Time (h)
B
inv1Δ tetON-SPinv1-inv2-YFP
-Tetracycline
+Tetracycline
GFP
DIC
C
Glucose
Sucrose
WT
inv1Δ
inv1-sfGFP

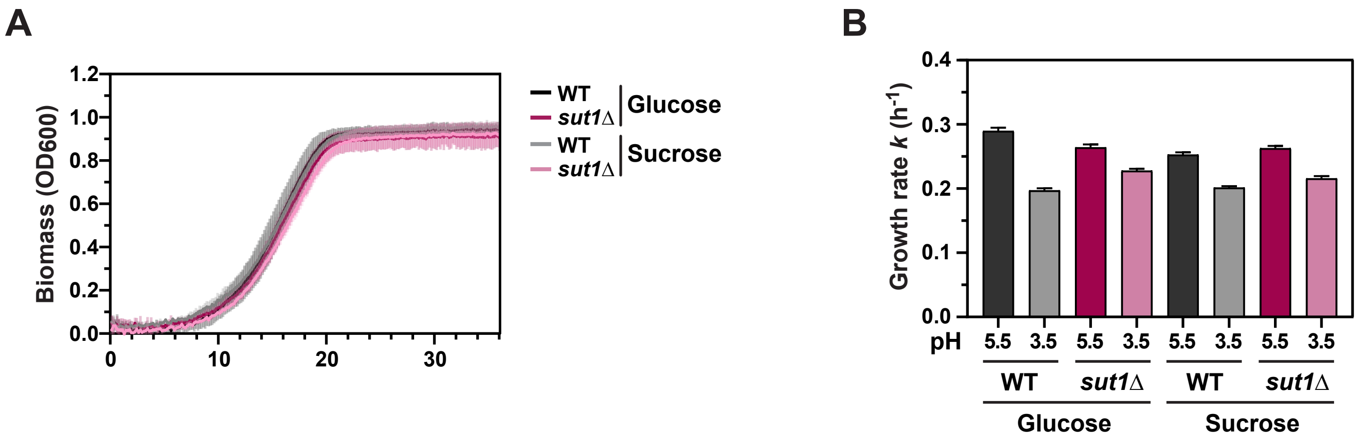
A
Biomass (OD600)
1.2
1.0
0.8
0.6
0.4
0.2
0.0
0
10
20
30
WT
sut1Δ
Glucose
WT
sut1Δ
Sucrose
B
Growth rate k (h-1)
0.4
0.3
0.2
0.1
0.0
pH
5.5
3.5
WT
sut1Δ
Glucose
Sucrose

**A**

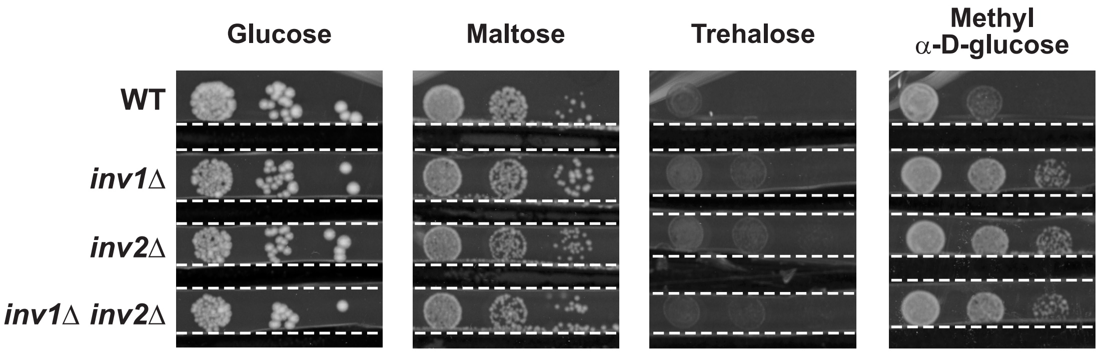


**B**

| | Glucose | Sucrose | Raffinose |
|---|---|---|---|
| WT | + | + | + |
| *inv1*Δ | + | - | + |
| *inv1*Δ *inv2*Δ | + | - | + |

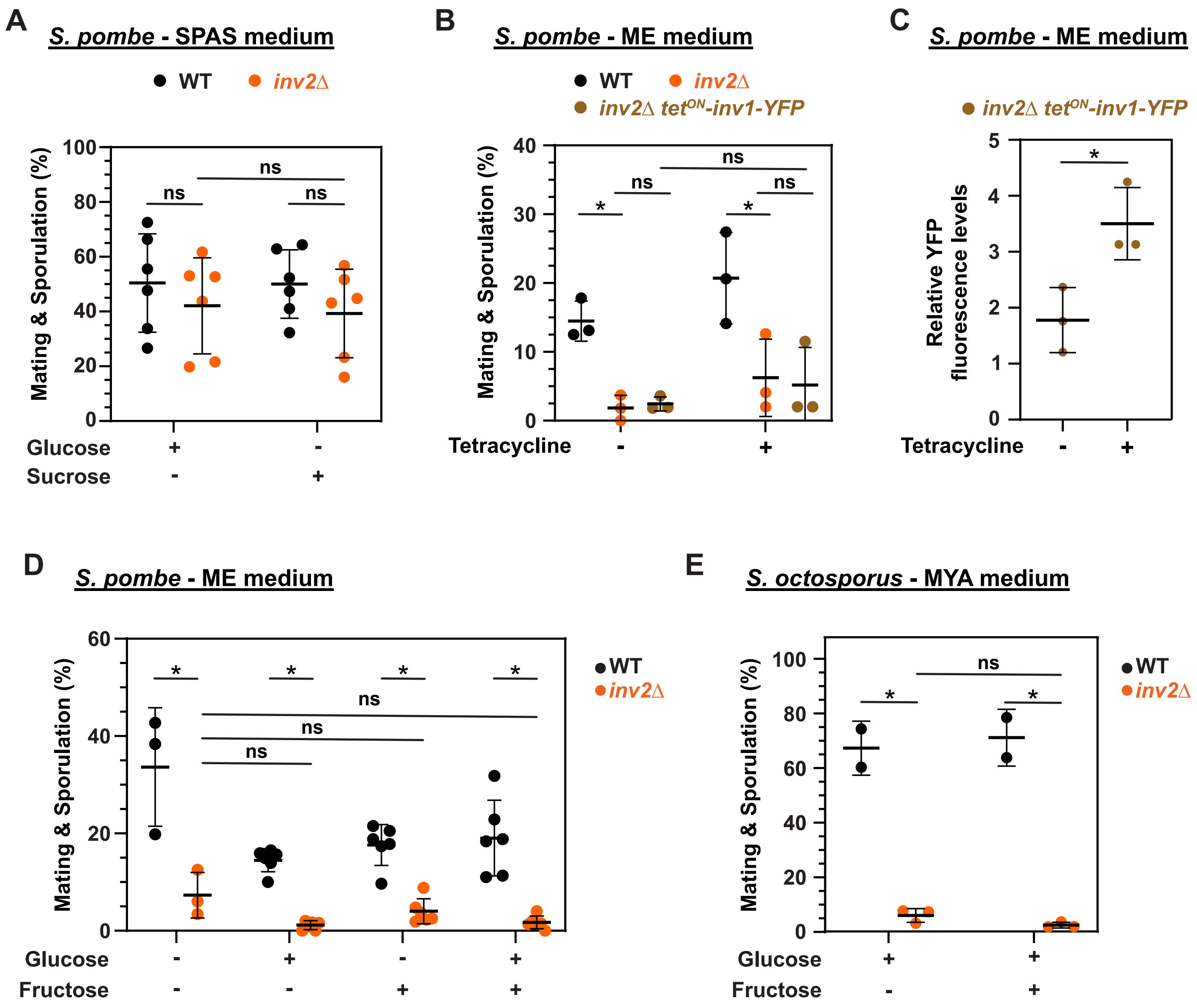
A
S. pombe - SPAS medium
WT
inv2Δ
Mating & Sporulation (%)
100
80
60
40
20
0
ns
ns
ns
Glucose + -
Sucrose - +
B
S. pombe - ME medium
WT
inv2Δ
inv2Δ tetON-inv1-YFP
Mating & Sporulation (%)
40
30
20
10
0
*
ns
ns
*
ns
Tetracycline - +
C
S. pombe - ME medium
inv2Δ tetON-inv1-YFP
Relative YFP fluorescence levels
5
4
3
2
1
0
*
Tetracycline - +
D
S. pombe - ME medium
WT
inv2Δ
Mating & Sporulation (%)
60
40
20
0
*
*
*
*
ns
ns
ns
Glucose - + - +
Fructose - - + +
E
S. octosporus - MYA medium
WT
inv2Δ
Mating & Sporulation (%)
100
80
60
40
20
0
ns
*
*
Glucose + +
Fructose - +

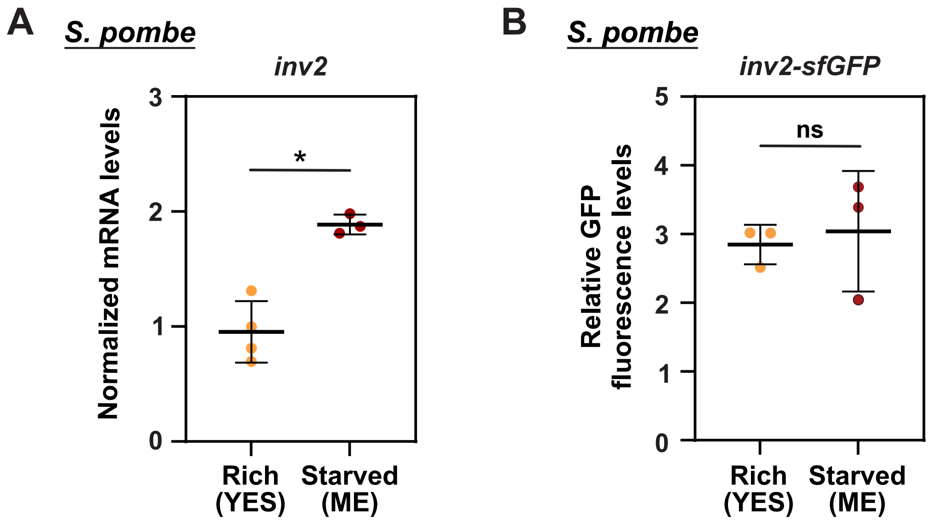
A
S. pombe
inv2
Normalized mRNA levels
3
2
1
0
*
Rich (YES)
Starved (ME)
B
S. pombe
inv2-sfGFP
Relative GFP fluorescence levels
5
4
3
2
1
0
ns
Rich (YES)
Starved (ME)
